# Protein-Peptide Turnover Profiling reveals wiring of phosphorylation during protein maturation

**DOI:** 10.1101/2022.04.03.486883

**Authors:** Henrik M. Hammarén, Eva-Maria Geissen, Clement Potel, Martin Beck, Mikhail M. Savitski

## Abstract

Post-translational modifications (PTMs) regulate various aspects of protein function, including degradation. Mass spectrometric methods that rely on pulsed metabolic labeling are very popular to quantify turnover rates on a proteome-wide scale. Such data have often been interpreted in the context of protein proteolytic stability. Here, we combine theoretical kinetic modeling with experimental pulsed stable isotope labeling of amino acids in cell culture (pSILAC) for the study of protein phosphorylation. We demonstrate that metabolic labeling combined with PTM-specific enrichment does not measure effects of PTMs on protein stability. Rather, it reveals the relative order of PTM addition and removal along a protein’s lifetime—a fundamentally different metric. We use this framework to identify temporal phosphorylation sites on cell cycle-specific factors and protein complex assembly intermediates. Our results open up an entirely new aspect in the study of PTMs, by tying them into the context of a protein’s lifetime.

## Introduction

Metabolic labeling coupled with mass spectrometry (MS) has become a mainstay of measuring protein turnover and degradation rates in cells. In cell culture experiments, arginine and lysine labeled with stable carbon and nitrogen isotopes are typically used in a method called pulsed (or dynamic) stable isotope labeling of amino acids in cell culture (pSILAC) (Mathieson et al., 2018; Schwanhäusser et al., 2011). Following labeling for a defined time, cells are lysed and proteins digested with trypsin, which cleaves after arginine and lysine residues, thus leaving each resulting peptide carrying at least a single labeled or unlabeled residue. Mass spectrometry is then used to measure the label incorporation rate for each identified peptide. In a steady-state system (or a steadily growing cell population, where the effect of cell growth can be accurately determined and subtracted from the data, see Theory Supplement), the incorporation rate equals the rate of clearance of the specific peptide from the system. In the case of an entire single protein species (or “proteoform”), which is cleared from the system by whole-protein degradation, the clearance rate equals the degradation rate. In this case, the degradation rate constant can be determined from the median clearance rate of all proteotypic peptides of a given protein (Claydon and Beynon, 2012; Mathieson et al., 2018; Schwanhäusser et al., 2011; Welle et al., 2016) (Figure 1).

**Figure 1:**
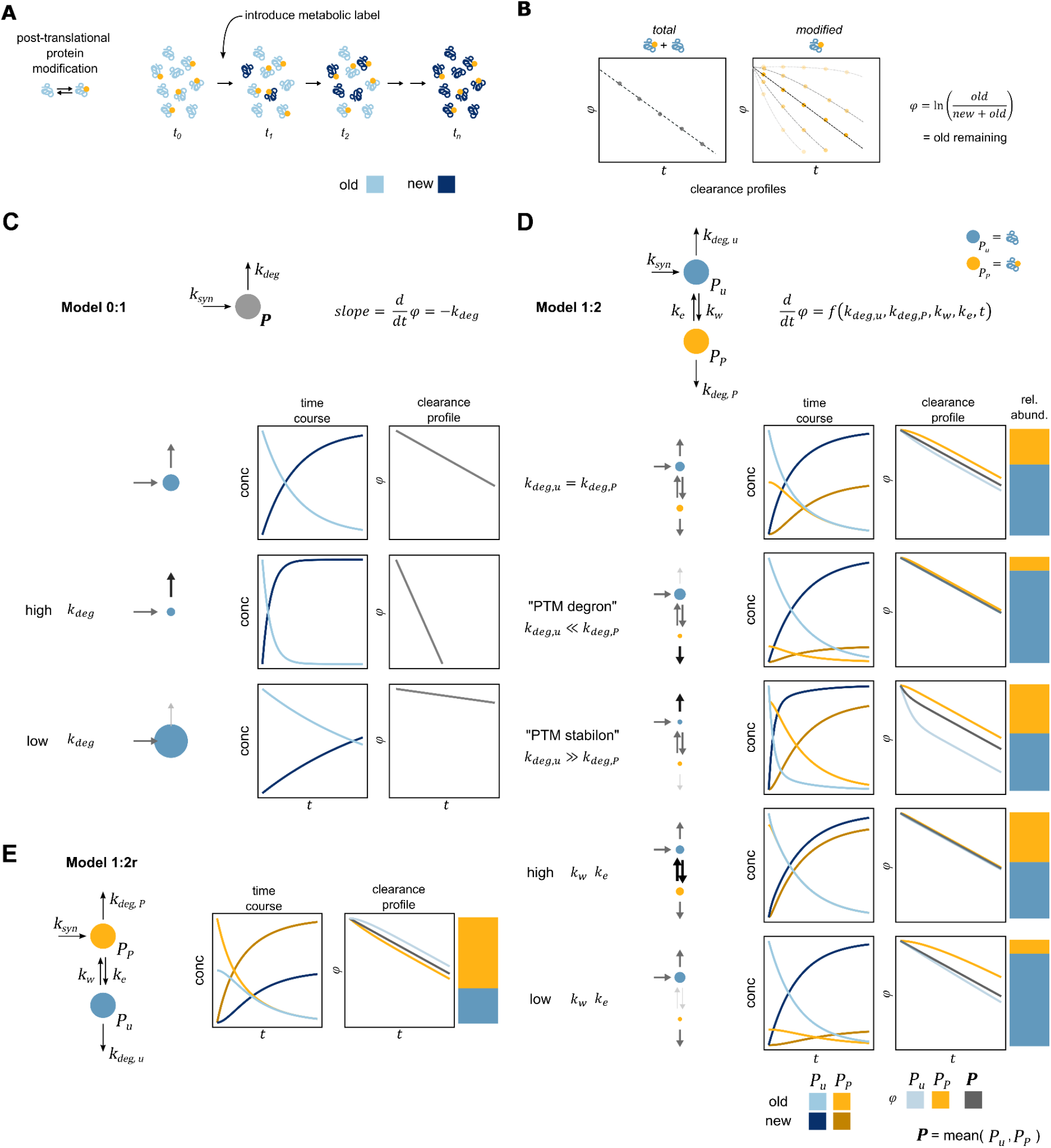
Quantitative modeling of multi-species metabolic labeling experiments identifies limits and possibilities of the approach. **A**, In a steady-state system with interconverting species, such as proteins with or without a PTM, an introduced metabolic label equilibrates into both the unmodified and the modified protein species through writing and erasing of the PTM (yellow circle). **B**, Hypothetical clearance profiles (logarithm of fraction of old protein remaining, *φ*, over time, *t*) for the entire protein and the modified species. **C**, In the simplest single-species model for protein turnover, the slope of the clearance profile is directly defined by the protein degradation rate constant, *k*_deg_. In the small cartoons next to the example traces, the size and color of the arrows reflect the size of the rate constant in question. The size of the circle reflects the steady-state amount of the protein species. **D**, Two-species model including protein modification, where the protein is synthesized in an unmodified form *P*_u_. The slope of the clearance profiles is a complex function of all model parameters (except the synthesis rate *k*_syn_), and changes over time (see Theory Supplement for full analytical description). In Model 1:2 clearance of the modified form *P*_P_ can never be faster than the clearance of the entire protein *P*. Also, parameter combinations with distinct biological interpretation, such as modifications causing rapid protein degradation (“PTM degrons”) or modifications with high interconversion rate constants (*k*_w_ and *k*_e_ for writing and erasing, respectively) have practically indistinguishable clearance profiles. **E**, Alternative two-species model, in which a protein is synthesized in a modified form, allows faster clearance of the modified species *P*_P_. Thus, the relative order of clearance profiles are defined by the order of species (or “wiring” of the modification network) during a protein’s lifetime. See also (https://hhammaren.shinyapps.io/PPToP_AllModels/) for an interactive web application of the models presented.

In reality, however, most eukaryotic proteins are present in more than a single proteoform. While some of these arise from alternative splicing of transcripts, the vast majority of cellular proteoforms is thought to be defined by post-translational modifications (PTMs) (Jensen, 2004), such as addition and removal of chemical groups (phosphorylation, acetylation, etc.) or proteolytic processing. These proteoforms often differ from one another only on a single amino acid residue (the one carrying the PTM), which can result in only a single proteoform-specific peptide after trypsin digest for MS analysis. The fate of these proteoforms is interlinked, as they are interconverted from one another, often in a reversible process of PTM writing and erasing. Crucially for metabolic labeling experiments, this interconversion can effectively “remove” a specific peptide from the system (by changing it to a differently modified peptide) by other means than degradation. As a consequence, the relation between clearance rate and degradation rate becomes nontrivial.

Here, we examine this relation using theoretical considerations based on first principles and experimentally test our resulting hypotheses in the context of protein phosphorylation. We show, using quantitative kinetic modeling, that clearance rates measured by pSILAC-MS for different proteoforms are not straightforwardly defined by effects of the PTMs on a protein’s proteolytic stability (i.e. the degradation rate). Rather, clearance rates and profiles are primarily defined by the network structure (or “wiring”) and affected by the kinetics of addition and removal of the modification itself. Thus, instead of readily yielding information on protein stability *per se*, differences in clearance rates allow deriving hypotheses on the temporal ordering of modification events along the synthesis-maturation-degradation axis (i.e. lifetime) of each protein. We test these hypotheses by combining pSILAC with phosphoenrichment and peptide-level turnover analysis in a method we dub Protein-Peptide Turnover Profiling (PPToP, similar to DeltaSILAC (Wu et al., 2021), or Site-resolved Protein Turnover profiling, SPOT (Zecha et al., 2022)). In accordance with our hypotheses, we find that the majority of phosphorylated peptides exhibit slower clearance than the respective protein median, characteristic of modifications occurring later in a protein’s lifetime. Furthermore, we identify peptides with faster clearance as expected, corresponding to known protein N-terminal maturation intermediate proteoforms. We corroborate our model by mutagenesis of 23 target proteins carrying 65 phosphosites. Using PPToP, we identify temporal proteoforms corresponding to phosphorylation/dephosphorylation events of cell cycle-specific factors as well as protein complex maturation intermediates.

## Results

### Proteoform clearance profiles in metabolic labeling are primarily defined by the proteoform modification network structure

In the case of a single pool of protein *P* in a steady-state system, the rate of clearance of the old protein (equal but opposite in sign to the rate of incorporation of new label) measured in a pSILAC-MS experiment is defined by a single parameter: the degradation rate constant of the protein, *k*_deg_ (Figure 1 B, C, E). To understand how adding a separate interconvertible protein pool (such as a proteoform defined by a PTM) affects measured clearance rates, we built quantitative kinetic models of synthesis-modification-degradation networks (Figure 1 D, G). The simplest model consists of two species: the unmodified protein, *P*_u_, and a modified protein, *P*_P_. Assuming a protein is synthesized in an unmodified form and is then modified in a first order reaction (see Theory Supplement for full descriptions and underlying assumptions), we define Model 1:2, where the rate of clearance of *P*_P_ and *P*_u_ become a function of not only the degradation constants of both protein species (*k*_deg,u_ and *k*_deg,P_), but also of the rate constants of interconversion (rate constants of writing and erasing the PTM, *k*_w_ and *k*_e_, respectively). Comparing the clearance rates of *P*_P_ and the entire protein pool *P* (= *P*_P_ + *P*_u_, measured in practice as the median clearance rate of all shared peptides), we find that *P*P exhibits slower clearance compared to *P* independently of the parameters used (Theory Supplement, and Figures 1 D and S1 A). Importantly, this not only includes cases, where a PTM might affect a protein’s proteolytic stability (so called “degrons”, when *k*_deg,P_ >> *k*_deg,u_, or “stabilons”, when *k*_deg,P_ << *k*_deg,u_), but also when there is no such effect. Thus, a slower clearance rate for *P*_P_ can not be interpreted to, for example, imply a stabilizing effect for the PTM in question. This stems from the fact that in Model 1:2 new label is introduced first into *P*_u_ (by synthesis) only from which it can subsequently enter into *P*_P_ through modification (Figure 1 D). Thus, the clearance rate of *P*P can at most equal the clearance rate of *P* as seen for rapid interconversion of the species (Figure 1 F: high *k*_w_ and *k*_e_). Furthermore, for cases, where the entire protein *P* has a linear clearance profile (which has been shown experimentally to be the case for most proteins (McShane et al., 2016), as well as constitutes the majority of theoretically achievable profiles), the magnitude of the difference in clearance of *P* and *P*_P_ is primarily defined by the writing and erasing rate constants (*k*_w_, *k*_e_, see Theory Supplement and Figure S1).

What does it take then, to produce faster clearance of *P*_P_ compared to *P*, as has been observed experimentally before (Wu et al., 2020; Zecha et al., 2022)? Arguably, the simplest solution is to relax the assumption that a protein is synthesized in an unmodified form, thus reversing the relative order of the two species (Model 1:2r, Figure 1 G). This can biologically be interpreted to represent modification during or immediately after translation. Analogously to Model 1:2, this causes clearance of *P*_P_ to always be faster or equal to *P*. Thus, relative clearance (or the “clearance profile”) is primarily defined by the relative order of appearance of the measured species with respect to protein synthesis, which is defined by the model structure, or “wiring”.

Naturally, networks of protein modification could be expanded to be arbitrarily complex to represent different biological situations. With increasing complexity of the model (and a resulting increase in parameters), the flexibility of the model increases concomitantly. Thus, for instance, a three-species model can be devised (Model 1:3, see Theory Supplement and Figure S7), where differences in clearance can also appear to be in line with differences in the degradation rate constants (*k*_deg,u_ and *k*_deg,P_). Most importantly, however, without prior knowledge of the shape of the network, these behaviors can not be unambiguously distinguished from effects caused by the relative order of modification along a protein’s lifetime using measured clearance rates of *P*_P_ and *P* alone (see Theory Supplement).

### Experimental detection of proteoforms with differing clearance rates by PPToP

The theoretical considerations above let us formulate three key predictions: (i) the default, (and thus functionally mostly uninformative) expected behavior for a PTM is slower clearance, (ii) interconvertible species with faster clearance likely represent early (i.e. close to synthesis) intermediates in protein maturation, and (iii) differences in relative clearance rates are not predictive of effects on protein proteolytic stability. To test these predictions for the case of protein phosphorylation, we devised PhosphoProtein-Peptide Turnover Profiling (Phospho-PPToP) combining two-label pulsed SILAC labeling with phosphoenrichment of peptides. We focused on early time points to enable observation of rapidly turning over transient species and sampled HeLa cells in 9 time points starting from 30 min after medium exchange (Figure 2 A). Following harsh lysis and trypsin digestion phosphopeptides were enriched, and both the phospho-enriched eluate and the total input were prefractionated and measured using data-dependent acquisition (DDA). We achieved high phosphoenrichment efficiency of 97%, and after filtering for reproducibility (Figure S2 A) and presence in at least 2 replicates and 2 time points, we could quantify 67393 unique modified peptides (covering 6749 gene names) carrying 10765 unique phosphosites (2880 gene names), including 2644 unique phosphosites (1065 gene names) for which peptides were quantified both in an unphosphorylated and phosphorylated form. Quantifying clearance of pre-existing (“old”) amino acids after medium exchange, we found that most unmodified peptides closely follow the protein median (Figure S2 B). This is expected, since most proteins should turn over as entire polypeptides, and only peptides typical to non-majority proteoforms should deviate from the median. Correspondingly, phosphopeptides exhibit much more varied clearance compared to the unmodified protein median with a tendency towards slower clearance (Figure S2 B), suggesting that the measured phosphorylated peptides indeed represent separate proteoform pools. The measured difference was not due to biases introduced in phosphoenrichment or measurement, as demonstrated by high correlation of clearance for peptides quantified in both the eluate and the total fraction (Figure S2 C).

**Figure 2:**
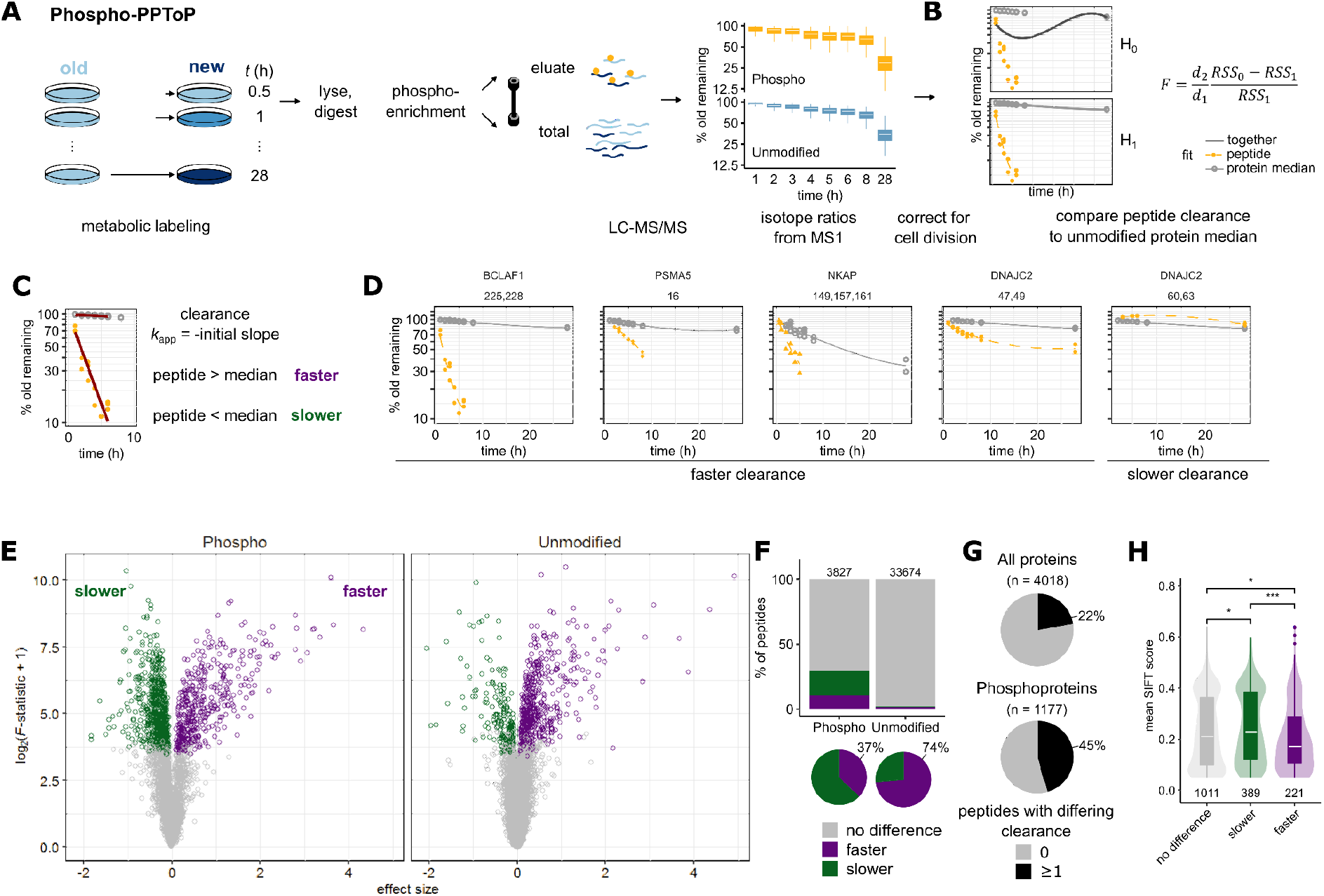
Measuring peptide clearance with PPToP. **A**, Experimental setup. HeLa cells were grown in isotopically Light (L)-labeled medium and pulsed for a specified time with Heavy (H)-labeled medium (Replicates 1–3; L and H reversed for Replicate 4). After lysis and trypsin-digest, phosphorylated peptides were enriched using iron-IMAC. Both the eluate enriched for phosphorylated peptides (phospho, in orange) as well as the flow-through (unmodified, in gray) were measured with mass spectrometry. Boxplots show accumulation of new labeled protein over time, y axis: median values over replicates on the protein level. **B**, After correction for cell growth, clearance traces for each peptide are compared to the trace of the median of all unmodified peptides (representing the total pool) to find peptides with significantly different clearance (see Methods for details). **C**, Example traces of phosphopeptides whose clearance significantly deviates from the protein median. Numbers over facets represent phosphosites phosphorylated on the peptides in question. **E**, Volcano-like plot showing peptides with significantly slower or faster clearance compared to the protein median. x axis shows the difference in fit compared to the protein median signed by the difference in initial slope: sign(Δinitial slope)*√(RSS_0_ - RSS_1_). **F**, Significantly more peptides in the phospho fraction exhibit differing clearance with the majority of hits being slower. In contrast, most hits of unmodified peptides exhibit faster clearance. **G**, Phosphoproteins are more likely to show differing clearance in at least one of their peptides compared to all proteins. **H**, Phosphosites exhibiting faster clearance are more likely to be conserved and functionally important as shown by lower SIFT scores (Dana et al., 2019; Ochoa et al., 2019).

We identified peptides with differing clearance using a comparative fitting approach comparing each peptide (unmodified or phosphorylated, representing *P*_u_ and *P*_P_, respectively) to the median of all (other) unmodified peptides of that protein, (representing the total protein pool, *P*) (Figure 2 B, see also Methods). To generate high-confidence hits, we only included peptides measured in at least 2 replicates, 4 time points, and with a minimum of 2 unique unmodified peptides to constitute the protein median reference. This resulted in statistical comparison for 3827 phosphopeptides (covering 1177 gene names), and 33673 unmodified peptides (3831 gene names). Hits were classified into “faster” or “slower” clearance based on the initial slope (≤ 6 h) of the clearance profile representing an initial clearance rate, *k*_app_ (Figure 2 C, D). In accordance with the larger overall spread of clearance of phosphorylated peptides, we found a significantly greater fraction of hits in phosphorylated than unmodified peptides (Figure 2 E, F), with 45% of all quantified phosphoproteins exhibiting at least a single peptide with significantly altered clearance (Figure 2 G). Hits were evenly distributed across phosphorylated amino acid residues (S, T, or Y, Figure S2 D) and protein abundance in both sample types (Figure S2 E).

It should be noted that, as a whole, our bottom-up approach is likely to underestimate differences in clearance rates of proteoforms, as measurements of peptides shared between multiple proteoforms represent an abundance-weighted mean over all proteoforms present (our *P*). Thankfully, however, this should only decrease the likelihood of false positives and increase the confidence of identified hits to actually represent distinct proteoform pools.

### Slower clearance of phosphopeptides is the expected default behavior and unlikely to be functionally informative alone

In line with our prediction from the simplest PTM-containing model (Model 1:2, Figure 1 D), we find the majority (63%) of phosphopeptide hits exhibiting slower clearance (Figure 2 F). Importantly, as demonstrated by Model 1:2, this is most readily explained by slow addition of phosphate groups (i.e. low *k*_w_). As this can result from numerous biological processes (including potential “non-functional” protein phosphorylation (Lienhard, 2008)), we deem these differences in clearance rate unlikely to be functionally informative when considered in isolation. This is also reflected in an overall younger evolutionary age of phosphosites with slower clearance (Figure S2 F), often equated with lower likelihood of functional importance (Ochoa et al., 2019).

Conversely, however, peptides with faster clearance demand further explanation and likely represent a functionally more interesting subgroup as also represented by lower SIFT scores corresponding to higher conservedness and higher predicted functional importance (Figure 2 H) (Dana et al., 2019; Ochoa et al., 2019).

### Faster clearance rates reveal protein maturation intermediates

In contrast to the slower clearance for the majority of phosphorylated peptides, we find that our unmodified hits are strongly enriched for faster clearance (74% of hits). Theory suggests that these peptides correspond to newly-synthesized or intermediate proteoforms, destined to undergo conversion to another form during protein maturation or aging (Model 1:2 and 1:2r, Figure 1 D, E). Indeed, this is what we observe, as unphosphorylated peptides with faster clearance are strongly enriched for peptides, where a corresponding phosphopeptide was also measured (Figure 3 A, B). Furthermore, the corresponding phosphopeptide showed predominantly slower clearance (Figure 3 C) suggesting directionality of the modification with regard to protein maturation in line with Model 1:2 (Figure 1 D). Similarly, peptides with faster clearance are also enriched for sites of other previously-identified PTMs (UniProt Consortium, 2021), (Figure 3 D). The same trend was observed for multiply-phosphorylated peptides, where the number of accumulated phosphate groups correlated with slower clearance (Figure 3 E, Figure S2 G), as expected from a stepwise modification cascade (Figure 3 F).

**Figure 3:**
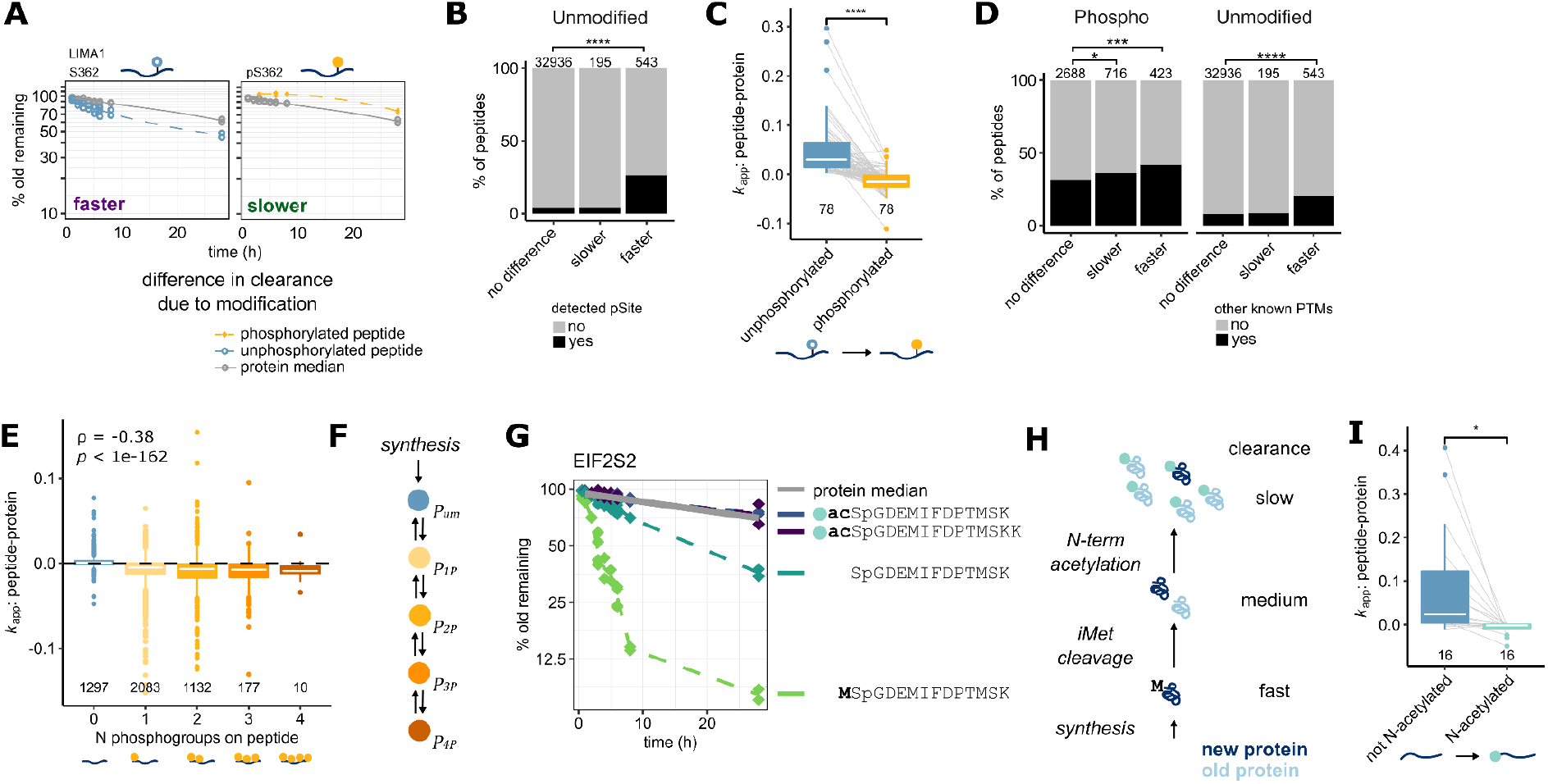
Peptide clearance rates reveal intermediates of protein modification. **A**, Change in clearance of a LIMA1 peptide caused by phosphorylation of Ser362. Left: faster clearance of LIMA1 peptides containing unmodified S362 (blue dashed line) compared to the median of the rest of the unmodified LIMA1 peptides (gray solid line). Right: slower clearance of LIMA1 peptides phosphorylated at S362 (orange dashed line). **B**, Unmodified peptides with faster clearance are strongly enriched for carrying amino acids also detected in a phosphorylated form (phosphosites, pSite). Fisher’s exact test, *p* < 2.2e-16. **C**, Initial clearance (k_app_) of peptides carrying pSites, where the unmodified form was identified as showing faster clearance and a phosphorylated form was also quantified. Shown are median values over replicates and peptides in case the same pSite was detected on multiple peptides. Paired *t* test, *p* = 2.6e-15. **D**, Phosphorylated peptides with faster and slower clearance, but only unmodified peptides with faster clearance are enriched for peptides carrying other known PTM sites (from Uniprot) suggesting that they represent early intermediates of protein modification cascades. Fisher’s exact test, **: *p* = 0.005; ****: *p* = 7.6e-14. **E**, Multiply-phosphorylated peptides trend towards lower clearance as number of phosphogroups increase. Peptides with zero phosphogroups are unmodified carrying residues, which were also detected in a phosphorylated form. Shown are peptides with slower or not significantly different clearance. ρ, Spearman correlation. These data suggest gradual, cumulative phosphorylation after synthesis as shown in **F**. **G**, Maturation intermediate peptides on the EIF2S2 N-terminus show faster clearance consistent with sequential, step-wise protein maturation depicted in the cartoon **H**. **I**, Non-acetylated N-terminal peptides from proteins undergoing N-acetylation have faster clearance consistent with them being maturation intermediates. Note that turnover of the final N-terminally acetylated form (in turquoise) closely follows the turnover rate of the protein median. Paired *t* test, *p* = 0.01.

Strikingly, the effect of sequential, directed modification on clearance is perhaps most clearly observable for N-terminal protein acetylation, a well-understood process thought to be irreversible in cells (Ree et al., 2018). We measured multiple N-terminal peptides of Eukaryotic Translation Initiation Factor 2 Subunit Beta (EIF2S2), whose relative clearance rates are concordant with stepwise, sequential maturation (Figure 3 G). The species with the fastest clearance still contains the initiator methionine (light green in Figure 3 G), which, once removed by N-terminal methionine excision, creates the second intermediate species (dark green in Figure 3 G). Finally, maturation is completed with N-terminal acetylation of the resulting peptide, whose clearance finally closely follows the median of all other EIF2S2 peptides, corresponding to the bulk of the fully mature EIF2S2 pool (Figure 3 G, H). In total, we found 16 examples of N-terminally acetylated proteins, where both an acetylated and an non-acetylated N-terminal peptide could be quantified (Figure 3 I; Figure S2 H, I), all of which showed relative clearance in line with our Model 1:2 for irreversible protein modification (Figure S2 J).

Additionally, our high-resolution dataset also includes more typical proteoform-level effects visible as differing clearance rates of a subset of peptides. These include, for example, isoform-specific turnover of Nuclear autoantigenic sperm protein (NASP), signs of autoproteolytic cleavage of Nucleoporin 98/96 precursor (Fontoura et al., 1999), and partial degradation of Nuclear Factor Kappa B Subunit 2 (NFKB2) (Sun, 2012) (Figure S3). However, these represent the minority of the peptide hits detected, and will thus not be discussed further in the context of this study.

### Fast clearance due to concerted dephosphorylation

The eukaryotic cell cycle is known to give rise to pervasive, synchronized, and ordered changes in both protein phosphorylation (Daub et al., 2008; Dephoure et al., 2008) and synthesis (Becher et al., 2018; Herr et al., 2020). Notably, this synchronization effect can be expected to remain visible in our PPToP data despite the use of asynchronous cultures, if phosphorylation and/or dephosphorylation occurs in a defined sequence with respect to a protein’s synthesis. A prime example of this is the proliferation marker protein Ki-67 (MKI67), for which we observe widespread faster clearance of phosphorylated peptides (Figure 4 A, Figure S4 B). Previous proteomic data (Herr et al., 2020) suggests that MKI67 is primarily synthesized in G2 phase coinciding with high MKI67 phosphorylation (Figure 4 B). Our data suggests that synthesis and phosphorylation is followed by an event of concerted MKI67 dephosphorylation. This could also explain the recently-described function of MKI67 in segregating premitotic chromosomes, which is followed by a molecular change in MKI67 leading to chromosome condensation (Cuylen-Haering et al., 2020).

**Figure 4:**
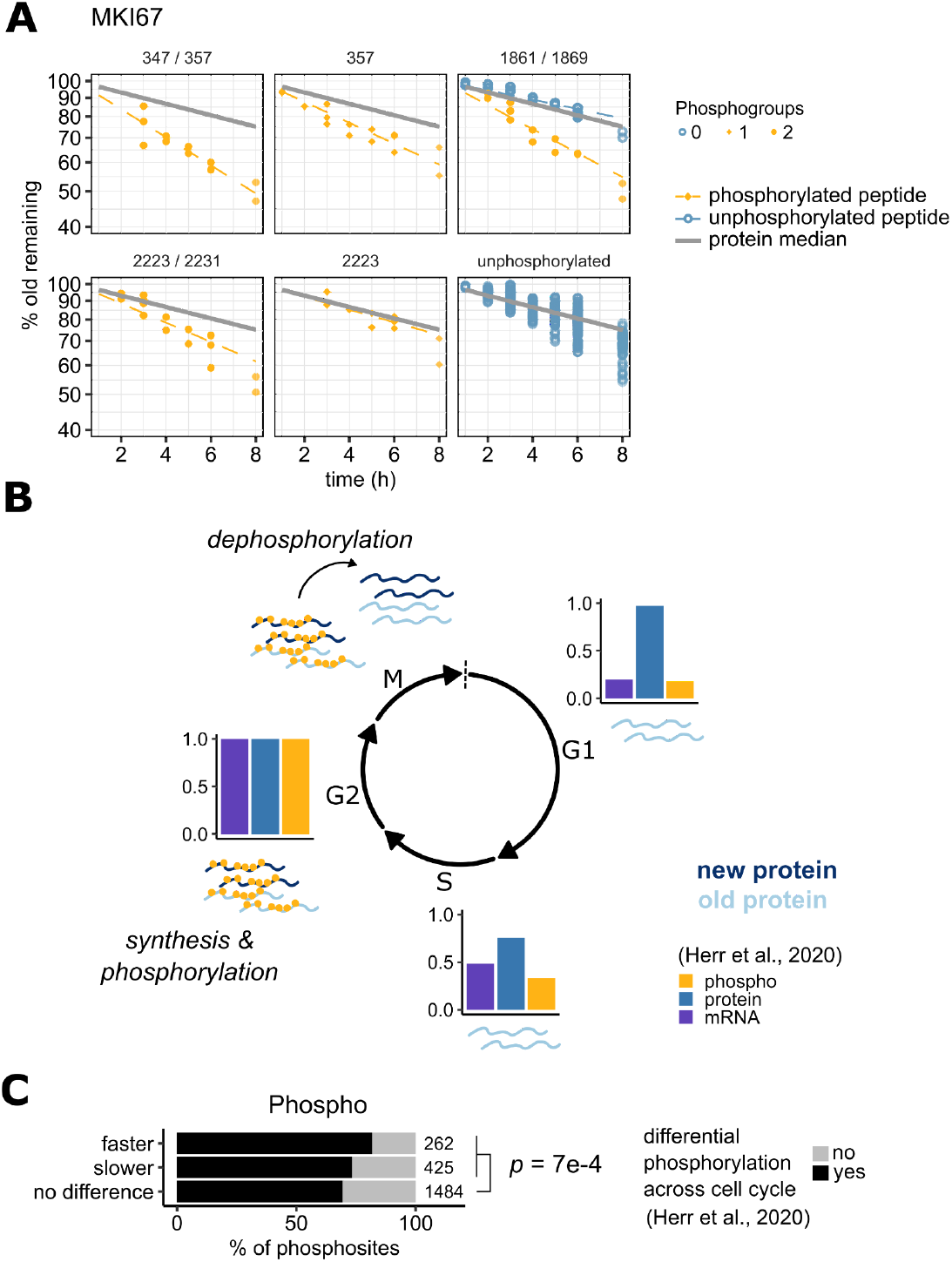
PPToP captures cell-cycle dependent effects. **A**, Representative examples of MKI67 phosphopeptides with faster clearance. Numbers above facets indicate phosphosites on the protein. Note that the peptide covering sites 1861 and 1869 was detected both phosphorylated (in yellow) and unphosphorylated (in blue) with distinct turnover profiles. **B**, Our data combined with previous data profiling the phosphoproteome across the cell cycle (Herr et al., 2020) suggest that MKI67 is synthesized and phosphorylated in G2 phase and subsequently undergoes concerted dephosphorylation in M phase. The dephosphorylation event suggested by the faster clearance measured by PPToP (A) explains previous reports of MKI67’s role in condensation of chromosomes during M phase (Cuylen-Haering et al., 2020). **C**, Hits from the phospho fraction are enriched for peptides exhibiting cell cycle-dependent changes in phosphorylation or abundance (Herr et al., 2020). Fisher’s exact test.

However, while proteins exhibiting cell cycle-dependent synthesis and/or phosphorylation patterns (such as MKI67) are slightly enriched in our PPToP dataset (Figure 4 C), they are unlikely to be the main contributing factor for the majority of hits found.

### Fast clearance is not predictive of low proteolytic stability in cells

Previous reports measuring turnover of PTM-modified peptides have focused on their potential effects on proteolytic stability, largely equating high measured clearance rates with fast protein degradation (Wu et al., 2020; Zecha et al., 2022), and interpreting PTM sites with fast clearance as potential degrons. In contrast, our theoretical considerations predict that measured changes in clearance can be unrelated to actual protein stability effects. We thus set out to test this prediction experimentally. We chose a diverse and representative set of 23 proteins carrying 65 phosphosites of interest (Figure 5 A) covering a wide range of protein half-lives. 53 out of the 65 chosen phosphosites showed fast clearance in our primary PPToP screen (Figure 5 A). We mutated each group of phosphosites to alanines (ALA, creating phosphorylation-incompatible) and/or aspartates (ASP, mimicking the negative charge of phosphoryl groups), and expressed them as mEGFP-fusions in HeLa cells (Figure 5 B) at roughly comparable expression levels (Figure S5 A). If the measured differences in clearance were due to differing proteolytic stability, one would expect the degradation rates of ASP mutants to correlate with the measured clearance rates, while ALA mutants should anti-correlate. We thus quantified their effects on the protein’s degradation rate by combining isobaric labeling and pulsed SILAC (Figure 5 B). Fusion proteins were pulled-down using anti-GFP beads under harsh buffer conditions to isolate them from endogenous proteins before digest. Comparing degradation rates of each mutant to the corresponding wild-type (Figure 5 C), we found that while most mutations (both ASP and ALA) slightly lowered degradation (i.e. slightly stabilizing the protein, Figure S5 B), these differences were not statistically significant for any of the sites tested (Figure 5 D). These data strongly suggest that differences in clearance measured by PPToP are independent of protein stability effects, and thus unlikely to directly represent degrons or stabilons.

**Figure 5:**
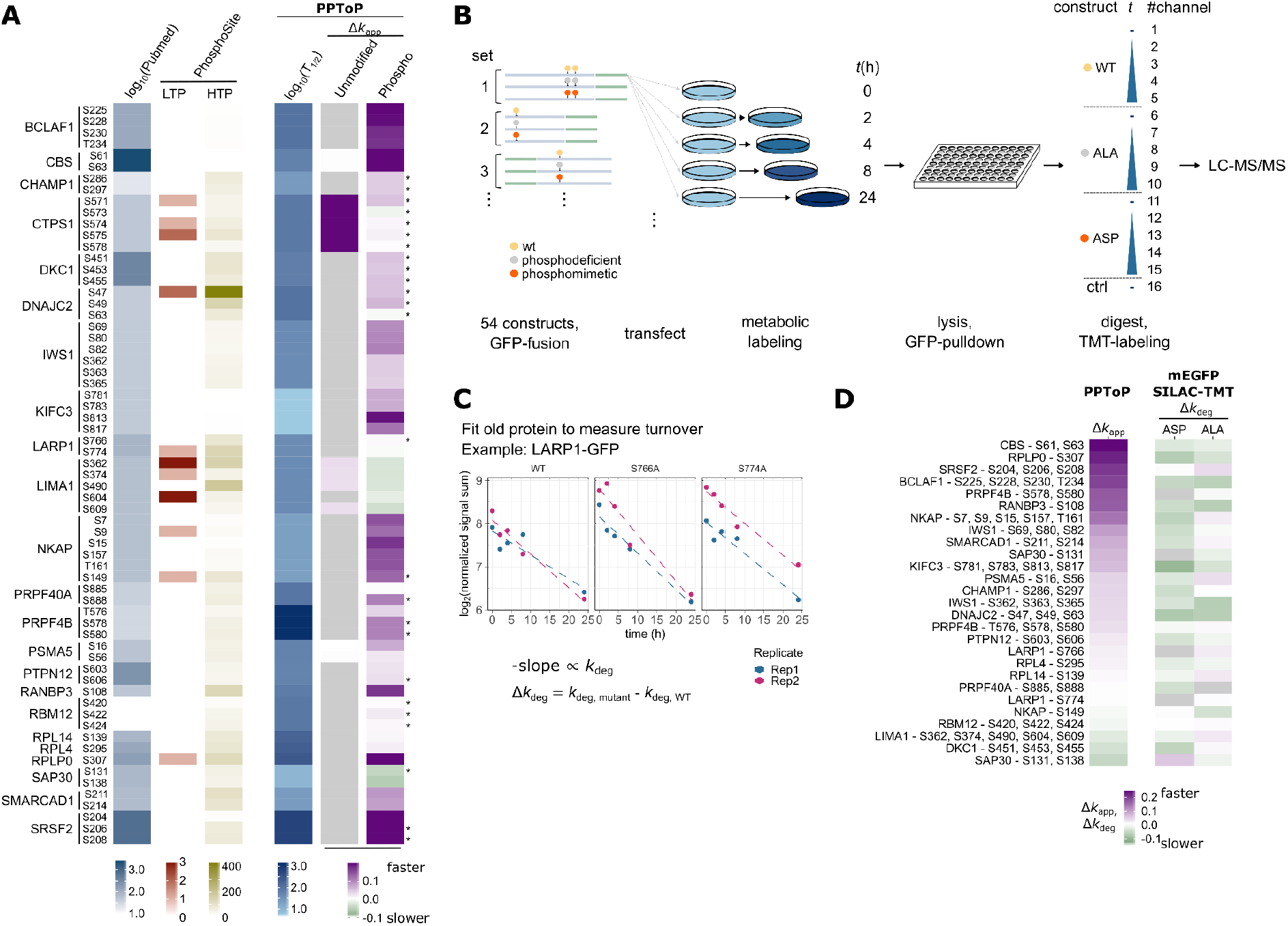
Differential clearance of phosphopeptides is not predictive of effects of phosphorylation on a protein’s proteolytic stability. **A**, Target phosphosites chosen for follow-up experiments. Representative targets were chosen to include a diverse mix of previous information available in the literature, as well as covering a wide range of protein half-lives (log_2_(T_1/2_) in hours, this study). Differential clearance of the unmodified or phosphorylated forms carrying the phosphosites of interest are shown. Cases, where the same phosphosite is present on multiple peptides with differing clearance rates are marked with asterisks. Cases where the unmodified form of the phosphosite was not quantified are shown in gray. Pubmed and PhosphoSitePlus were accessed on 16.11.21 and searched using the gene name or common alternative names, where applicable. LTP = Low-throughput, HTP = high-throughput reference studies. **B**, Experimental setup for interrogating effects of phosphosite mutations on protein proteolytic stability. **C**, Turnover of exogenously expressed proteins was quantified from TMT signal sums of the old protein from pull-down experiments. **D**, Comparison of differences in clearance rates from PPToP (Δ*k*_app_) and wild-type-to-mutant differences from the exogenous expression experiment show no significant correlation (see also Figure S5 B), indicating that differences in clearance measured from PPToP are not predictive of differences in protein degradation caused by the PTM.

### Maturation intermediate phosphosites can affect protein-protein interactions and protein complex assembly

As PPToP can detect protein maturation intermediates, we looked at protein complex assembly, where control of maturation is especially important as protein-protein interactions need to be established in a controlled manner, and unwanted interactions could be highly deleterious. Interestingly, our dataset includes phosphorylation sites with fast clearance on known protein complex subunits, representing prime candidates for maturation intermediates. In particular, we followed-up two phosphorylation sites: S16 and S56 on Proteasome subunit alpha type-5 (PSMA5). When phosphorylated, both sites exhibit faster clearance than their corresponding unphosphorylated peptides, which closely match the protein median (Figure 6 A), suggestive of transient early phosphorylation (Model 1:2r). Interestingly, both sites lie on protein-protein interfaces with neighboring subunits (PSMA1 and PSMA7) in the intact proteasome alpha ring and neither is phosphorylated in the mature complex (Figure 6 B). Pull-downs with wild-type (WT) PSMA5-GFP under stringent buffer conditions showed reproducible cosedimentation of PSMA1, while neither the phosphodeficient (ALA) nor the phosphomimetic (ASP) mutant was able to pull down PSMA1 (Figure 6 C). Additionally, previous profiling of thermal stability effects of phosphorylation also showed these sites to be associated with significantly lowered thermal stability of PSMA5 in HeLa cells (Potel et al., 2021) (Figure 6 D) as well as in yeast (Smith et al., 2021), suggesting that phosphorylated PSMA5 is not part of the assembled proteasome, as proteins assembled into larger complexes tend to show higher thermal stability (Mateus et al., 2018; Tan et al., 2018). Taken together, these data suggest that interaction with PSMA1 relies on functional phosphorylation of PSMA5 (most likely at S16), and that the phosphorylated species is not associated with the intact proteasome complex. Based on our PPToP data we suggest that phosphorylated PSMA5 represents a protein complex assembly intermediate potentially required for correct folding of PSMA5 (Figure 6 E).

**Figure 6:**
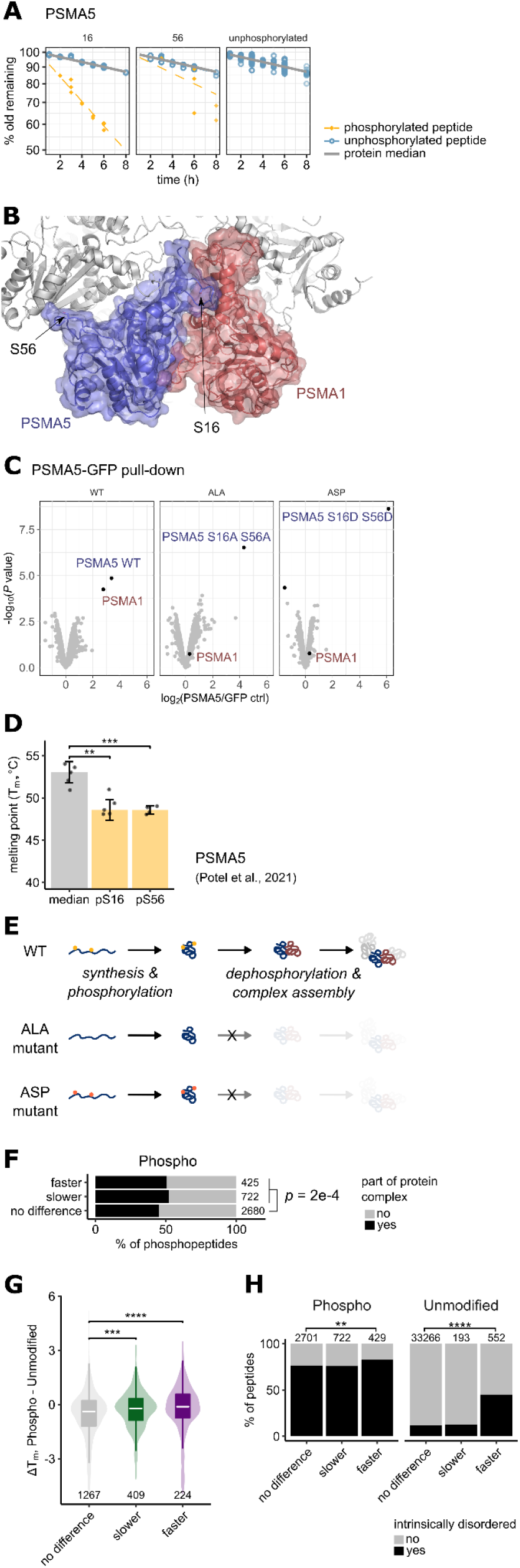
Transient phosphorylation at S16 on PSMA5 is needed for interaction with PSMA1. **A**, PPToP traces of PSMA5 peptides carrying S16 or S56, as well as all unphosphorylated peptides are shown. **B**, Structure of PSMA5 (in blue) in the mature proteasome. The neighboring alpha ring subunit PSMA1 is highlighted in red. S16 and S56 are highlighted as sticks. Both residues are unphosphorylated in the mature proteasome structure (PDB ID: 6MSB). **C**, Pull-down under stringent buffer conditions of exogenously expressed PSMA5-GFP constructs in HeLa cells shows that the PSMA5-PSMA1 interaction is lost upon mutation of S16 and S56. Shown is the unadjusted *P* value from a limma analysis. N = 2. **D**, Thermal stability of peptides phosphorylated at S16 and S56 is significantly lower than the PSMA5 median. Data from (Potel et al., 2021). *t* test, **: *p* < 0.01; ***: *p* < 0.001. **E**, Model of the role of S16 (and S56) phosphorylation. We hypothesize that transient S16 and S56 phosphorylation is required for PSMA5 maturation and its incorporation into the proteasome. **F**, Phosphopeptides exhibiting differing clearance are enriched in protein complex subunits. Fisher’s exact test. Protein complexes are from the CORUM core complex dataset (Giurgiu et al., 2019). **G**, Phosphopeptides with differing clearance exhibit altered thermal stability compared to the unmodified protein median suggesting altered molecular states such as protein-protein interactions. Proteome-wide thermal stability data from (Potel et al., 2021). Wilcoxon test, ***: *p* < 0.001, ****: *p* < 1e-4. Outliers have been omitted for clarity. **H**, Peptides with faster clearance are enriched in intrinsically-disordered protein regions. Disorder prediction from D2P2 (Oates et al., 2013). See Materials and Methods for details.

We find that phosphorylated peptides with differing clearance are modestly, but significantly enriched in protein complex subunits also globally (Figure 6 F). Furthermore, comparing with earlier thermal proteomic profiling data (Potel et al., 2021), we find that differing clearance of phosphopeptides is also associated with changed thermal stability (Figure 6 G), again suggesting a distinct molecular state *in cellulo* for proteoforms defined by peptides identified by PPToP. We also find that peptides with faster clearance are enriched in intrinsically disordered regions of proteins (Figure 6 H), suggesting that these protein regions might act as specific switches between proteoforms, and that controlling the behavior of these regions (e.g. with PTMs) might be of special importance in the context of protein maturation. The enrichment in disordered regions in peptides with faster clearance was significant also after controlling for the confounding variable of other known PTMs, which tend to also be enriched in disordered regions (Figure S4 C). Furthermore, our follow-up data also includes examples, where mutations of phosphosites identified by PPToP control the tight interactome of the protein, such as association of SRSF2 with the RNA splicing machinery (Figure S6).

## Discussion

PTMs, such as reversible phosphorylation, control all aspects of protein function from protein-protein interactions and catalytic activity to subcellular localization and proteolytic stability. Despite advances in the identification and localization of PTMs onto proteins by MS-based omics technologies, functional annotation and understanding of PTMs is severely lagging behind as, e.g., >95% of known human protein phosphosites lack any annotation on biological function (Needham et al., 2019). This lack of understanding of PTMs is especially pronounced in the temporal dimension. Specifically, how (or whether) PTMs are added at specific times over a protein’s lifetime from its synthesis on a ribosome, folding, maturation and function, through to its eventual degradation, has so far remained largely elusive.

Based on experimentally validated theoretical considerations, we show here that peptide-level turnover analysis, such as PPToP, can be used to deliver exactly this information. Analogously to isotopic labeling in metabolic flux experiments (Jang et al., 2018), combining SILAC with MS effectively reveals the relative temporal order of events through observation of label incorporation along a network. Similarly, while we focus mainly on proteoforms defined by the addition and removal of PTMs, the same considerations and conclusions apply to any analogous metabolic labeling experiment of biomolecules in which a species can exist in multiple measurable states that can interconvert, such as nucleic acids and their modifications (Gameiro et al., 2021).

The information provided by PPToP allows establishing temporal ordering of events along the protein’s lifetime. It thus complements previous findings, which have estimated that for around 10% of human proteins degradation rates are dependent on the age of the protein itself (McShane et al., 2016). Our findings extend this notion by not only providing the opportunity to generate hypotheses about the PTMs involved in age-dependent stabilization, but also showing how wide-spread protein age-dependent modification is in the human proteome.

Using PPToP we identify numerous phosphosites of interest, including sites on PSMA5, which we think could correspond to proteasome assembly intermediates. Interestingly, we do also observe effects of previously known proteasomal protein maturation events in our data. Multiple proteasomal β subunits undergo proteolytic processing during proteasome assembly (Murata et al., 2009), including PSMB7, which is cleaved after residue 43. Our PPToP data includes a glimpse of this, as we detected a PSMB7 peptide starting at residue 42 with very fast clearance, despite the peptide only being detected in two time points (see PSMB7 in the interactive data browser accessible at https://hhammaren.shinyapps.io/PPToP_Data_browser/).

Recently, two groups have published analyses of pSILAC data with the goal of achieving proteoform-resolution of protein turnover by estimating the turnover of single, proteoform-specific peptides (Zecha et al., 2022), and this approach has been proposed to reveal the effects of PTMs on protein stability (Wu et al., 2021). Using insight from our theoretical modeling, we show here that the majority of differences in measured clearance rates are not linked to protein proteolytic stability differences, but rather are indicative of the rate of PTM addition and network wiring. Results by Zecha et al. (Zecha et al., 2022) on lysine acetylation showing preferentially slower rates of clearance are thus suggestive of slow rates of addition, which is in line with the general very low stoichiometry of lysine acetylation (Narita et al., 2019).

We also directly test the hypothesis that proteoform clearance rates are indicative of proteolytic stability. Using a substantial number of representative target proteins in our mutagenesis experiment, we found no positive correlation between differences in clearance rates from PPToP and differences in cellular turnover rates of our phosphosite mutants and wild-type constructs (Figure 5 D, Figure S5 B). In fact, we found a slight negative correlation for the phosphomimetic aspartate mutant (the opposite what would be expected for actual phospho-degrons), but even though this was marginally statistically significant, the magnitude of the effect was very small (Figure S5 B) and thus unlikely to be biologically significant. Based on both our theoretical predictions and our experimental validation, we thus conclude that proteoform clearance rates are not directly indicative of proteolytic stability effects.

Interestingly, pSILAC analysis of ubiquitinated peptides has shown a propensity for higher rates of clearance for peptides carrying ubiquitin-remnant sites (Zecha et al., 2022). This is noteworthy, as it suggests that these ubiquitination events occur preferentially on relatively recently-synthesized proteins, compared to the rest of the protein’s proteoforms. Faster clearance rates were especially enriched for ubiquitinated ribosomal proteins, and we speculate that many of these identified ubiquitination sites might act as quality control switches during ribosome biogenesis, potentially marking misfolded or misincorporated subunits that might be subsequently degraded. This behavior is in line, e.g., with our Model 1:3 (Figure S7), where fast clearance for the modified pool can be combined with rapid degradation in the case of an early bifurcation of protein fate (Figure S7 B: “early degron”). Notably, however, the same behavior can also be explained by deubiquitination during protein maturation, which would likewise lead to faster clearance rates (Figure S7 B: “maturation intermediate PTM site”). More targeted experiments, including mutagenesis of the identified sites, as well as careful examination of the clearance profile of the total protein profile (McShane et al., 2016) will be needed to distinguish between these two potential explanations.

In summary, we found that peptides exhibiting faster clearance are enriched, and thus are likely explained, by at least the following mechanisms: other PTMs (leading to further modification and concurrent clearance of the original peptide), concerted synthesis and (de)phosphorylation along the cell cycle, and protein maturation during complex formation. Interestingly, however, a large proportion of peptides with faster clearance currently remain without immediate mechanistic explanation (Figure S8). We think these previously unannotated sites thus present an exciting source for future research.

## Acknowledgements

We thank Dr. Sindhuja Sridharan for valuable discussions and critical discussions regarding the manuscript, Dr. Nils Kurzawa for assistance with the comparative F-statistic approach, Maria Zimmermann-Kogadeeva for helpful discussions on model analysis, Cecilé Le Sueur for fruitful discussions regarding various statistical analyses, the EMBL Proteomics Core Facility, particularly Mandy Rettel, for excellent technical assistance with the mass spectrometry measurements and instrumentation, as well as all members of the Savitski and Beck teams for feedback and input.

This work was supported by the European Molecular Biology Laboratory. H.M.H., and C.M.P. were supported by a fellowship from the EMBL Interdisciplinary Postdoctoral (EI3POD) Programme under Marie Skłodowska-Curie Actions COFUND (grant number 664726).

## Author contributions

Conceptualization, H.M.H. and M.M.S.; Visualization, Data Curation, and Investigation H.M.H.; Formal Analysis, H.M.H and E.-M.G.; Mathematical Analysis of Models, E.-M.G.; Methodology, H.M.H., C.M.P.; Writing - Original Draft, H.M.H. and E.-M.G.; Writing - Review & Editing; H.M.H., E.-M.G., M.B., and M.M.S.; Project Administration, H.M.H.; Resources and Supervision, M.B. and M.M.S.

## Materials and Methods

### Cell culture and isotopic labeling

HeLa Kyoto cells (from S. Narumiya, RRID: CVCL_1922) were cultured according to standard tissue culture techniques in SILAC DMEM Flex Media (Gibco) containing either “Light” (^12^C and ^14^N-labeled, Arg0 and Lys0, Thermo) or “Heavy” (^13^C and ^15^N-labeled, Arg10 and Lys8, Silantes) labeled arginine and lysine, 1 mg/ml glucose, 10% dialyzed FBS (Gibco), and 1 mM L-glutamine. Cells were grown for a minimum of 10 doublings in the respective media before starting a time course to ensure complete labelling. For the SILAC time course 2.3E6 cells were seeded onto 15 cm dishes in Light (replicates 1 through 3) or Heavy medium (replicate 4), cultured for 24 h to 48 h before medium exchange. A single 15 cm dish was used per time point per replicate. Medium was aspirated and cells washed twice with warm PBS (with Ca and Mg), then medium replaced with Heavy (replicates 1 through 3) or Light medium (replicate 4). For each replicate all dishes were seeded and lysed simultaneously with staggered medium exchange.

Both cell lines used were verified to be mycoplasma free.

### Sample preparation and phosphoenrichment

Cells were washed twice with ice-cold PBS and scraped on ice into lysis buffer (6M Urea, 100 mM Hepes (pH8.5), 5 mM Tris(2-carboxyethyl)phosphine, 30 mM chloroacetamide, 4 mM MgCl2, 2 mM NaVO4, 2 mM NaF, 2 mM Na-pyrophosphate, 1% sodium deoxycholate, 1% Triton X-100, 1x Complete protease inhibitor cocktail (Roche), 1x PhosStop phosphatase inhibitor cocktail (Roche), 0.5% Benzonase (Merck, 70746)), diluted 1:1 with dilution buffer (6M Urea, 100 mM Hepes (pH8.5), 4 mM MgCl2), and sonicated at +4°C in a Bioruptor Plus, 45 cycles, 30 s on / 30 s off (Diagenode). Lysates were cleared by centrifugation, 16 000 g, at 4°C for 60 min and supernatants frozen to −80 °C before continuing. Cleared supernatants were thawed at room temperature (RT) and nucleic acids digested by adding 0.5% fresh Benzonase and incubating 1 h at RT, after which EDTA was raised to 25 mM and SDS to 2%. Protein was precipitated in a 4:4:1 lysate:methanol:chloroform mixture, centrifuged 10 min at 4000 g, RT, and the resulting protein precipitate extracted and washed twice with 70% EtOH in a sonicator water bath. Protein was resuspended to 2.5 mg/ml into digestion buffer (100 mM Hepes (pH8.5), 5 mM Tris(2-carboxyethyl)phosphine, 30 mM chloroacetamide, 1% sodium deoxycholate), TPCK-Trypsin (Thermo, 20233) added to 100 μg/ml and incubated on an end-over-end shaker overnight at RT. Digested peptides were desalted on SepPak columns using gravity flow (Waters, WAT054945), washed twice with 0.1% trifluoroacetic acid (TFA), eluted with 40% acetonitrile (ACN), and resulting peptides lyophilized.

Phosphopeptides were enriched essentially as previously described (Potel et al., 2018, 2021) using a ProPac Immobilized Metal Ion Affinity Chromatography (IMAC)-10 column (Thermo, 063276) loaded with Fe3^+^ on a Dionex Ultimate 3000 HPLC system. The phosphopeptide-containing eluate as well as the flow-through were collected and lyophilized.

#### High-pH peptide prefractionation

5% of the flow-through (“total”) was taken up in 20 mM ammonium formate (pH 10) and prefractionated first into 29 fractions on a 1200 Infinity HPLC (Agilent) using high-pH reversed-phase chromatography (running buffer A: 20 mM ammonium formate pH 10; elution buffer B: ACN) on an X-bridge column (2.1 × 10 mm, C18, 3.5 μm, Waters). Fractions were then pooled across to generate 12 fractions and vacuum dried.

Phosphopeptides were fractionated manually using in-house packed C18 microcolumns as previously described (Potel et al., 2021) into seven fractions.

### Exogenous expression of GFP-fusion constructs, pull-down, and sample preparation

23 proteins identified as carrying phosphorylation sites of interest from the proteome-wide experiment were cloned as either N or C terminal fusions to monomeric EGFP (carrying the A206K mutation) in a vector under a Ubc promoter (kind gift from Daniel Heid and Judith Zaugg). Phosphorylation sites of interest were mutated to either alanine and/or aspartate (see Table SN for full list of constructs). Cloning and mutagenesis was performed by GenScript (GenScript Biotech Netherlands B.V.).

Constructs were overexpressed in HeLa cells as follows. Cells were seeded onto 12-well plates at 90E3 cells per well in Light SILAC medium the day prior to transfection using a single well per construct, per time point, per replicate. Each well was transfected with 360 ng of plasmid using Lipofectamine 3000 (Invitrogen) according to manufacturer’s instructions, and let to transfect for 24 h, after which medium was exchanged to Heavy medium for all wells simultaneously. Cells were collected after the designated time in Heavy medium, placed on ice, washed with PBS and lysed in lysis buffer (50 mM Hepes pH7.5, 150 mM NaCl, 0.5 mM EDTA, 0.1% SDS, 1% Triton X-100, 1% deoxycholate). After lysis, cells were collected by scraping, diluted 1:1 with dilution buffer (50 mM Hepes pH7.5, 150 mM NaCl, 5 mM MgCl_2_, 0.625 U/ul Benzonase (Merck, 70746)), and left on ice for >30 min. Lysates were cleared by spinning 5 min at 2000 g at 4 °C, and filtering through a pre-wetted 0.22 μm filter plate (Merck millipore). For the GFP pull-down, cleared lysate was incubated with 2 μl washed GFP-Trap magnetic agarose beads (Chromotek) per well for >4 h at 4 °C, washed twice with wash buffer with NP40 (50 mM Hepes pH7.5, 150 mM NaCl, 5 mM MgCl_2_, 0.05% NP40), and thrice with detergent-free wash buffer (50 mM Hepes pH7.5, 150 mM NaCl, 5 mM MgCl2). Proteins were digested on beads with trypsin and Lys-C (5 ng/μl final concentration each) in 90 mM HEPES (pH 8.5), 5 mM chloroacetic acid and 1.25 mM TCEP overnight at room temperature shaking at 500 rpm. Peptides were eluted using 2% DMSO and dried in a speedvac.

Dry peptides were reconstituted in 5 μl water and labeled by adding 2 μl TMT label (20 μg/μl in acetonitrile (ACN)) (TMTpro 16 plex, Thermo Fisher Scientific) and incubating 1 h at room temperature. Labeling was quenched with hydroxylamine (1.1% final concentration), and samples pooled to make full TMT16 sets as shown in Figure 5B. Pooled sets were desalted on an OASIS HLB μElution plate (Waters 186001828BA); washing thrice with 0.05% FA, eluting with 80% ACN, 0.05% FA, and drying in a speedvac.

### Mass spectrometry

#### Proteome-wide PPToP

For LC-MS/MS analysis, peptides were reconstituted in 0.1% FA, 4% ACN and analyzed by nanoLC-MS/MS on an Ultimate 3000 RSLC (Thermo Fisher Scientific) connected to a Fusion Lumos Tribrid (Thermo Fisher Scientific) mass spectrometer, using an Acclaim C18 PepMap 100 trapping cartridge (5μm, 300 μm i.d. x 5 mm, 100 Å) (Thermo Fisher Scientific) and a nanoEase M/Z HSS C18 T3 (100Å, 1.8 μm, 75 μm x 250 mm) analytical column (Waters). Solvent A: aqueous 0.1% FA; Solvent B: 0.1% FA in ACN (all solvents LC-MS grade from Fisher Scientific).

LC-MS/MS analysis parameters for total proteome and phosphofraction were as follows: Peptides were loaded on the trapping cartridge using solvent A for 4 min (3 min for phospho) with a flow of 30 μl/min. Peptides were separated on the analytical column at 40 °C with a constant flow of 0.3 μl/min applying a 100 min gradient of 4–25% of solvent B in A, followed by a 5 min gradient (25–40%), and a 4 min washing step at 85% solvent B (both total and phospho). Peptides were directly analyzed in positive ion mode with a spray voltage of 2.4 kV and an ion transfer tube temperature of 275 °C (both total and phospho). Full scan MS spectra with a mass range of 300–1500 m/z (375–1500 m/z for phospho) were acquired on the orbitrap using a resolution of 120 000 (60 000 for phospho) with a maximum injection time of 50 ms (20 ms for phospho) and Normalized AGC Target of 50% (Standard for phospho). Data-dependent acquisition was used with a cycle time of 2 s (3 s for phospho). Precursors were isolated on the quadrupole with an intensity threshold of 1e3 (2e5 for phospho), charge state filter of 2–7, and an isolation window of 1.2 m/z (1.4 m/z for phospho). Precursors were fragmented using HCD at 30% (32% for phospho) collision energy. For the total proteome: MS/MS spectra were acquired on the ion trap, with a maximum injection time of 50 ms, and a dynamic exclusion window of 45 s. For the phosphoenriched fraction, MS/MS spectra were acquired on the orbitrap, at 30 000 resolution, a maximum injection time of 75 ms, and a dynamic exclusion window of 20 s.

#### AP-MS for analysis of mEGFP-fusion proteins

TMT16-labeled peptides from pull-down experiments were reconstituted in 0.1% FA, 4% ACN and analyzed by nanoLC-MS/MS on the same hardware described above. LC-MS/MS analysis parameters were as follows: Peptides were loaded on the trapping cartridge using solvent A for 3 min with a flow of 30 μl/min. Peptides were separated on the analytical column at 40 °C with a constant flow of 0.3 μl/min applying a 104 min gradient of 6–28% of solvent B in A, followed by a 4 min gradient (28–40%), and a 4 min washing step at 80% solvent B. Peptides were directly analyzed in positive ion mode with a spray voltage of 2.2 kV and a ion transfer tube temperature of 275 °C.

Full scan MS spectra with a mass range of 375–1500 m/z were acquired on the orbitrap using a resolution of 120 000 with a maximum injection time of 50 ms. Data-dependent acquisition was used in top 10 mode. Precursors were isolated on the quadrupole with an intensity threshold of 2e5, charge state filter of 2–7, and an isolation window of 0.7 m/z. Precursors were fragmented using HCD at 34% collision energy. MS/MS spectra were acquired on the orbitrap, at 30 000 resolution, a maximum injection time of 100 ms, scan range in first mass mode (at 110 m/z), and a dynamic exclusion window of 20 s.

#### Data availability

The mass spectrometry proteomics data will be deposited to the ProteomeXchange Consortium via the PRIDE (Perez-Riverol et al., 2022) partner repository.

### Data analysis

All data analysis was carried out in R (version 4.0.0 or later) (R Core Team, 2020).

#### Proteome-wide PPToP

##### Analysis of MS raw files

For PPToP MS raw files were processed using MaxQuant (version 1.6.4.0) (Cox and Mann, 2008) using a reference human proteome (uniprot Proteome ID: UP000005640, downloaded 9.6.2020). Data was processed separately for total and phosphoenriched samples, but for each all time points, fractions, and replicates were run together. Default search parameters were used, except as follows: multiplicity: 2; Heavy channel: Arg10, Lys8; variable modifications: Acetyl (Protein N-term), Oxidation (M), and only for the phosphofraction: Phospho (STY); fixed modifications: Carbamidomethyl (C); maximum number of modifications per peptide: 5; maximum missed cleavage sites: 2 (3 for phospho); LFQ: none; re-quantify: unchecked; match between run: checked.

##### Data filtering and preprocessing

Identified peptides from the MaxQuant evidence file were filtered to remove hits from the reverse database and potential contaminants. All subsequent analysis was done on the modified peptide level (henceforth referred to as “peptide”) including information on Heavy amino acid incorporation, N-terminal acetylation, and phosphorylation, but excluding methionine oxidation. Peptides quantified in only one of the SILAC channels (constituting 45.3% of all identified peptides) were removed. In case a peptide was quantified multiple times, a single entry was chosen by choosing the species with (1) the lower posterior error probability (PEP), and (2) the highest intensity. Peptides were further filtered for presence in at least 2 replicates and 2 time points.

Cell cycle times for each replicate were estimated from the protein median values in the unmodified fraction. Assuming exponential decay of most proteins, we have the linear relationship (Schwanhäusser et al., 2011):

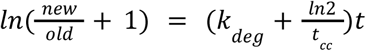

where 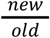 is the SILAC ratio of new and old protein, *k_deg_* is the protein-specific degradation constant, and *t_cc_* the cell cycle time. We estimated *t_cc_* from the 1% longest-lived proteins where, assuming no active protein degradation (*k* = 0), the slope is defined by 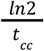, from which we got *t_cc_* estimates of 28.0 h, 26.5 h, 27.0 h, and 22.2 h for replicates 1 through 4, respectively. Subsequently, new/old SILAC ratios were transformed into fraction of old remaining (see Theory Supplement for reasons of doing so), and corrected for cell cycle as follows:

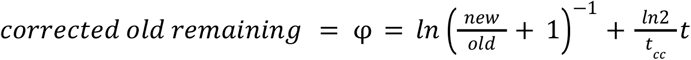

Next, peptide entries were filtered for reproducibility between replicates by calculating the distance of the corrected old remaining (φ) value for each peptide for each replicate to the median value of that peptide at that time point over all replicates, and excluding entries with values deviating from the median by more than two standard deviations of the entire distance distribution of all peptides at all time points in that fraction (unmodified or phospho). This removed 2.3% of all measurements.

##### Comparative fitting to find peptides deviating in clearance from the rest of the protein

The clearance of each peptide was compared to the median of all other quantified peptides of that protein from the unmodified fraction by fitting a spline with 3 degrees of freedom to the trace of *φ* vs time. Peptides with data in at least 4 time points, a total of at least 6 data points, and for which the median of the other peptides in that protein included at least 2 unique peptides in the unmodified fraction were included. An *F-*statistic was calculated for fitting the spline to either both peptide and protein median together (H0 model) or to each separately (H1 model). Due to heteroscedasticity of the data, the resulting *F*-statistic was calibrated as delineated in (Childs et al., 2019) by estimating the “effective degrees of freedom” (*d*_1_, *d*_2_) from a null dataset, where “peptide” or “median” labels were randomized (thus reducing any differences between the two classes to those occurring by chance). Since our dynamic PTM-SILAC dataset had differing amounts of data points per case, *d*_1_, *d*_2_ were estimated for six separate bins of the data depending on the number of data points available for the comparison. The thusly calibrated *F*-statistic distribution across both fractions was used to calculate *p* values for each peptide using the `pf ()` function from the *stats* package in R and corrected for multiple testing using Benjamini-Hochberg correction. Cases with adjusted *p* value <= 0.001 were considered as hits.

##### Disordered protein prediction

Prediction of protein disorder was taken from the D2P2 database (Oates et al., 2013) using a consensus threshold of 75% across the individual predictor algorithms when determining the disorder status per amino acid. For the enrichment analysis, a peptide was considered intrinsically disordered if it contained at least 40% disordered amino acids.

##### Other statistical analyses

Statistical tests were done using R using the following functions: Fisher’s exact test: `fisher.test()`; *t* test and Wilcoxon signed-rank test: `stat_compare_means()`; linear regression: `lm()`.

#### Targeted analysis of exogenously expressed GFP-fusion proteins

##### Analysis of MS raw files

MS raw files were processed using IsobarQuant (Franken et al., 2015) and peptide and protein identification was obtained with Mascot 2.5.1 (Matrix Science) using a reference human proteome (uniprot Proteome ID: UP000005640, downloaded 9.6.2020) modified to include the overexpressed protein constructs in question, known common contaminants and reversed protein sequences. Mascot searches were done only for old (light SILAC label) proteins. Parameters were: Trypsin/P; max. 3 missed cleavages; peptide tolerance 10 ppm; MS/MS tolerance 0.02 Da; fixed modifications: Carbamidomethyl (C), TMT16plex (K); variable modifications: Acetyl (Protein N-term), Oxidation (M), Phospho (ST), Phospho (Y), TMT16plex (N-term).

##### Turnover analysis of exogenously expressed proteins

Data from pull-down experiments was analyzed from the IsobarQuant peptide output file, filtering peptides to exclude contaminants (including skin and keratin contaminants), peptides without reporter (TMT) quantification data, peptides lacking K or R residues (e.g. C-terminal peptides), and peptides with FDRs > 0.01. In cases where a modified peptide was measured multiple times, the entries were collapsed to a single value choosing the peptide with the highest score and highest precursor-to-threshold (p2t) values. As the vast majority of peptides in all pull-downs were shared, and data was to be analyzed comparing each mutant construct to the respective, in-set WT sample, reporter signals were median-normalized over all channels. Next, construct-specific peptides were identified and excluded from the analysis in channels, where the construct in question was not experimentally expressed (e.g., peptides carrying an ALA mutation were excluded from quantification in channels containing WT and ASP expressing samples and vice versa). This was done to prevent TMT-induced bleed-through affecting the quantification. Constructs with low expression levels (<20% of the median expression level of all constructs in that set) were also removed, as well as peptides mapping to GFP, since some constructs also produced free GFP as seen on immunoblots. To fit degradation constants (*k*_deg_) signal sums of the remaining peptides of each construct were calculated and fitted with a linear fit over the linear portion of the time course (the last time point was removed for constructs with rapid degradation).

To estimate statistically, whether there are differences in degradation rates, fold changes of the signal sum of each mutant over the corresponding wild-type (within each set) were calculated for each time point. Linear fits on fold changes against time were done using the `lm()` function in R and resulting probabilities of non-zero slope (Pr_t) (indicating, whether there’s a difference in slope between the mutant and the corresponding wild-type) were corrected for multiple testing using the Benjamini-Hochberg method in the function `p.adjust ()`.

##### Co-pull-down interactome analysis

Changes in the tight interactome of the mutants compared to wild-type were estimated from the Isobarquant protein output file at time point zero (before label switch). Briefly, proteins were filtered as described above, and signal sum values were normalized to a GFP only channel in each set. For SRSF2, the interactome was visualized using the STRING plugin in Cytoscape (v 3.7.1). For PSMA5, differences in interactions were verified in a separate pull-down experiment without SILAC pulse in triplicate, and differences in interaction were identified by applying a limma analysis (Ritchie et al., 2015) on the fold changes.

## Supplemental information

This manuscript contains a separate Theory Supplement presenting the analysis of the quantitative protein synthesis-modification-degradation models.

The quantitative models and their behavior in a pSILAC experiment can be accessed via an interactive web application (https://hhammaren.shinyapps.io/PPToP_AllModels/)

The experimental data presented in this study can also be browsed and visualized via an interactive web application (http://hhammaren.shinyapps.io/PPToP_Data_browser/)

### Supplemental figure legends

**Figure S1:**
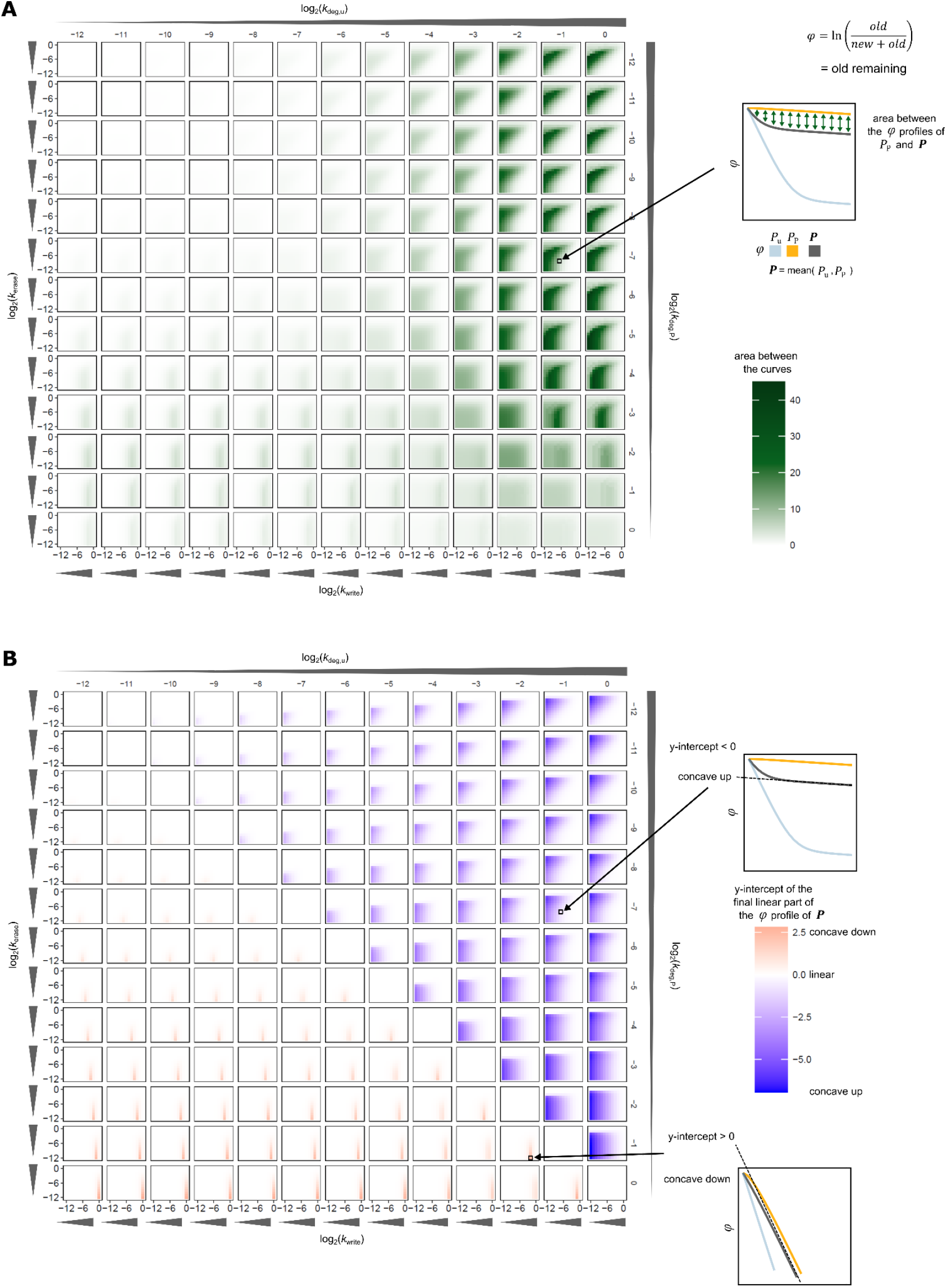
Example output of Model 1:2 with different parameter combinations. **A**, Sum of the distance between the clearance profile of the modified species, *P*_P_, and the entire protein *P* for a simulation of 0-24 h. Darker colors reflect larger differences in clearance profile. Note that the distance is always positive, as the modified species always exhibits slower or similar clearance than total (Model 1:2). For calculating the difference, values of *φ* corresponding to less than ~0.3% old remaining were omitted as smaller ratios are not practically measurable with pSILAC-MS. **B**, Same parameter combinations as in A, but showing the shape of the total clearance profile by plotting the extrapolated y-intercept of the linear end part of the curve. Nonlinear concave up curves yield a negative y-intercept (blue), while concave down curves give positive y-intercepts (red). Note that while the largest differences in clearance (A) can be achieved when *k*_deg,u_ >> *k*_deg,P_, this parameter combination simultaneously leads to a strongly nonlinear, concave up clearance curve (B) for the entire pool *P. k*_syn_ was not varied and is not shown, as it does not affect the clearance profiles.

**Figure S2:**
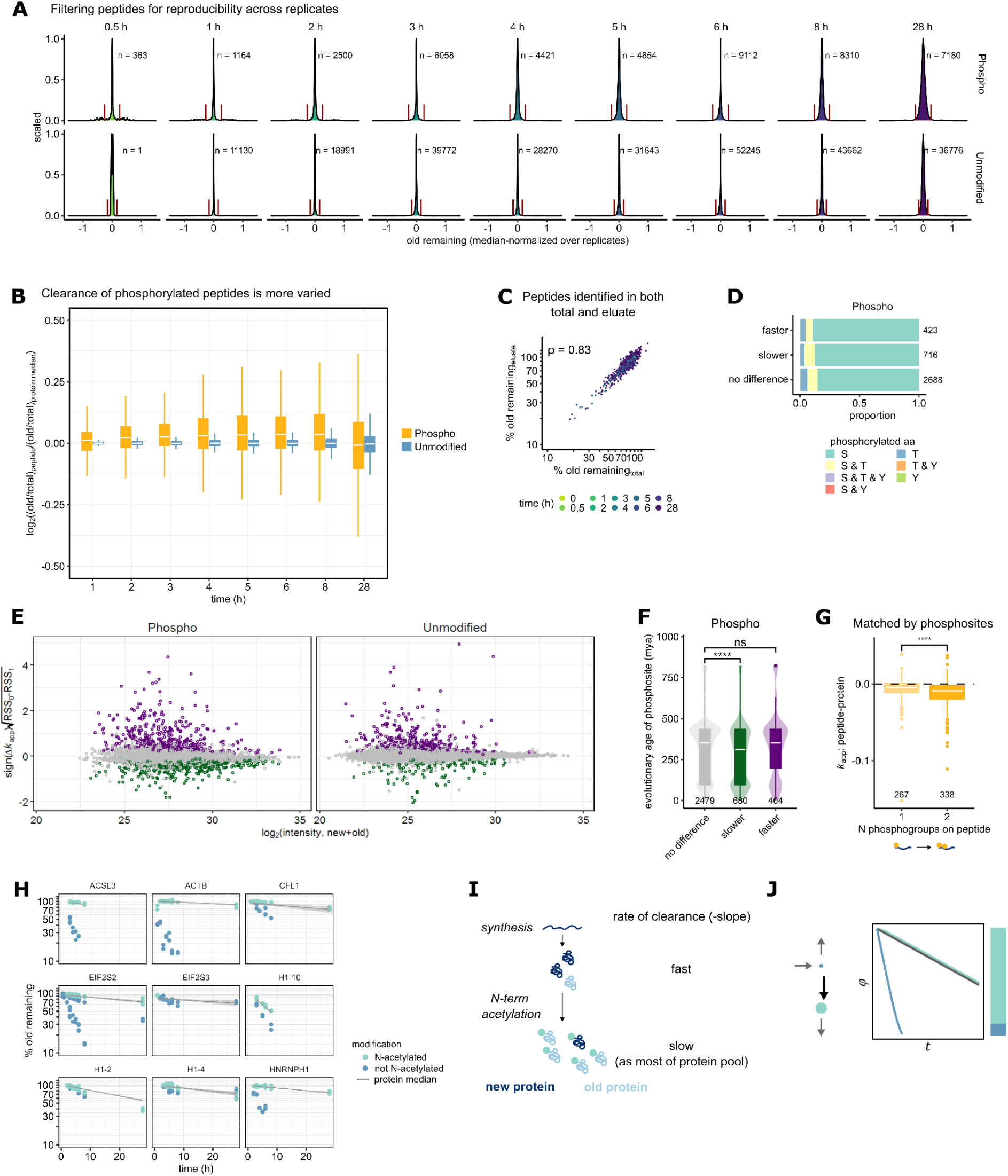
Experimental measurement of clearance rates with PPToP. **A**, Filtering of clearance values (from SILAC ratios) for reproducibility across replicates. **B**, Phosphopeptides show overall slower, but more varied clearance than unmodified peptides. Medians over replicates are shown. Outliers are omitted for clarity. **C,** High reproducibility of clearance measured for peptides quantified in both eluate and flow-through of the phosphoenrichment procedure shows that enrichment does not introduce artifacts in measured SILAC ratios. Shown are clearance values corrected for cell cycle and filtered for reproducibility across replicates. ρ, Spearman correlation. **D**, Hits from PPToP are evenly distributed across different phosphorylated residues, as well (**E**) as across peptide abundance. **F**, Phosphopeptide hits with slower clearance are evolutionarily younger than the rest of phosphosites. Data from (Ochoa et al., 2019). **G**, Addition of second phosphate group causes decrease in clearance rate. Related to Figure 3 E. Shown are all peptides which include exactly 2 detected phosphosites within their boundaries, and which are phosphorylated either once or twice. Peptides detected to have faster clearance were omitted as in Figure 3 E. ****, p = 2.4e-6, unpaired t-test. Including also peptides with significantly faster clearance does not change the direction, nor significance of the effect (p = 2e-5, data not shown). **H**, Representative plots of proteins undergoing N-terminal acetylation. Non-acetylated N-terminal peptides are shown in blue and the corresponding acetylated forms shown in teal. The median clearance of the protein is shown as a gray line. **I**, Cartoon of the linear protein maturation cascade of irreversible protein N-terminal acetylation. **J**, Model 1:2 showing the theoretical clearance profile for the unmodified species (in blue), the modified species (in teal) as well as the total protein (in gray) for rapid, irreversible protein modification, such as N-terminal acetylation.

**Figure S3:**
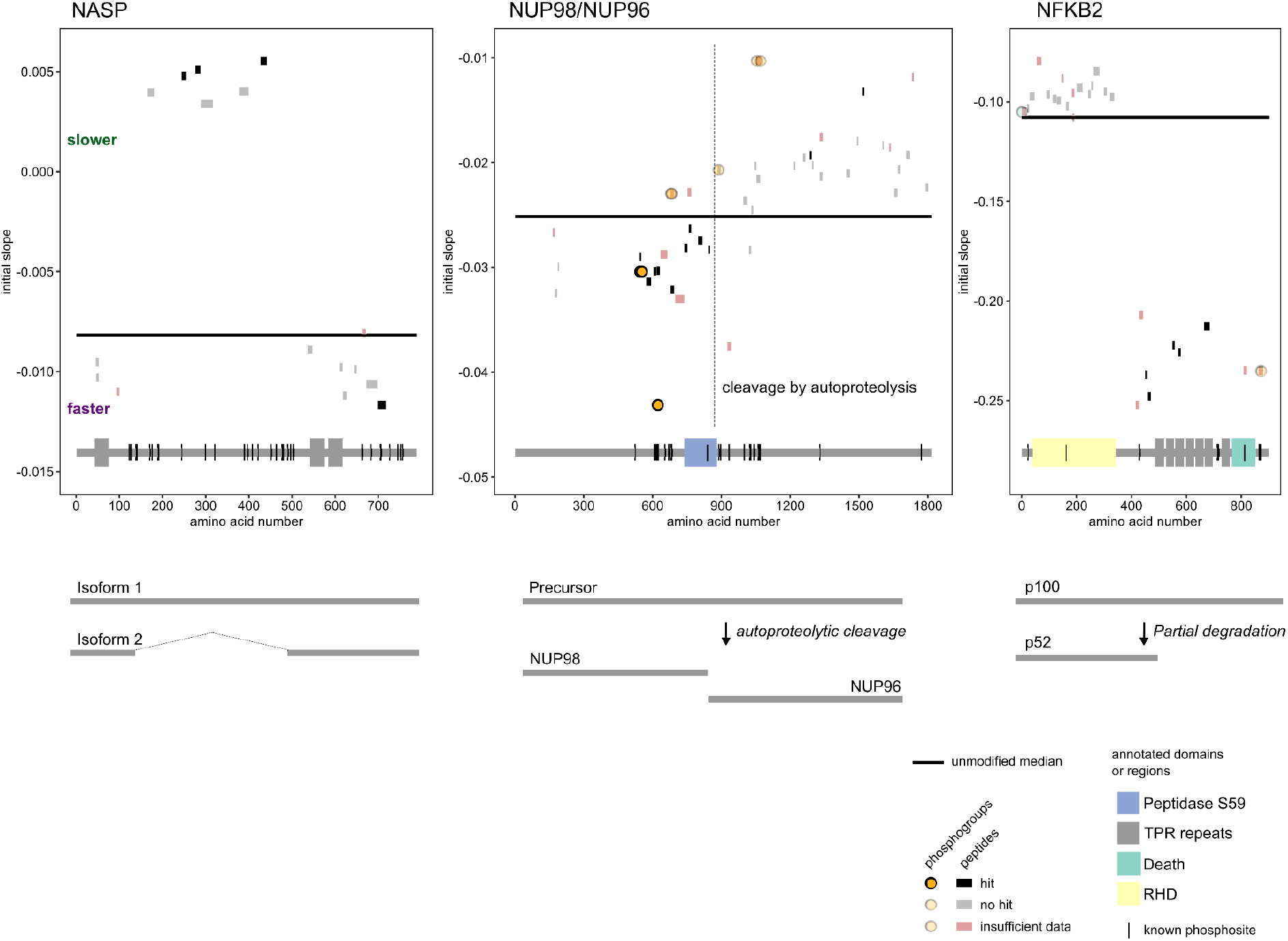
Protein-level turnover effects uncovered by PPToP. Three examples of intra-protein differential clearance rates due to protein-level effects such as alternative splicing generating isoforms with differing turnover (Nuclear autoantigenic sperm protein, NASP, left panel), post-translational protein cleavage to generate two proteins from a single precursor (Nucleoporins 96 and 98, NUP96/98, middle panel), or partial protein degradation generating a signaling competent proteoform (Nuclear factor NF-kappa-B, NFKB2, right panel). Each line segment (black or grey) shown represents a unique modified peptide detected along the protein’s sequence. The median initial clearance rate (*k*_app_) over all replicates is shown on the y axis for each peptide. The median initial clearance rate fitted to all unmodified peptides is shown as a dark line. Hits from the comparative fitting approach are indicated. Detected phosphosites are shown with spheres. Underneath is a cartoon depiction of the protein’s primary sequence with annotated domains and known phosphorylation sites (from Uniprot).

**Figure S4:**
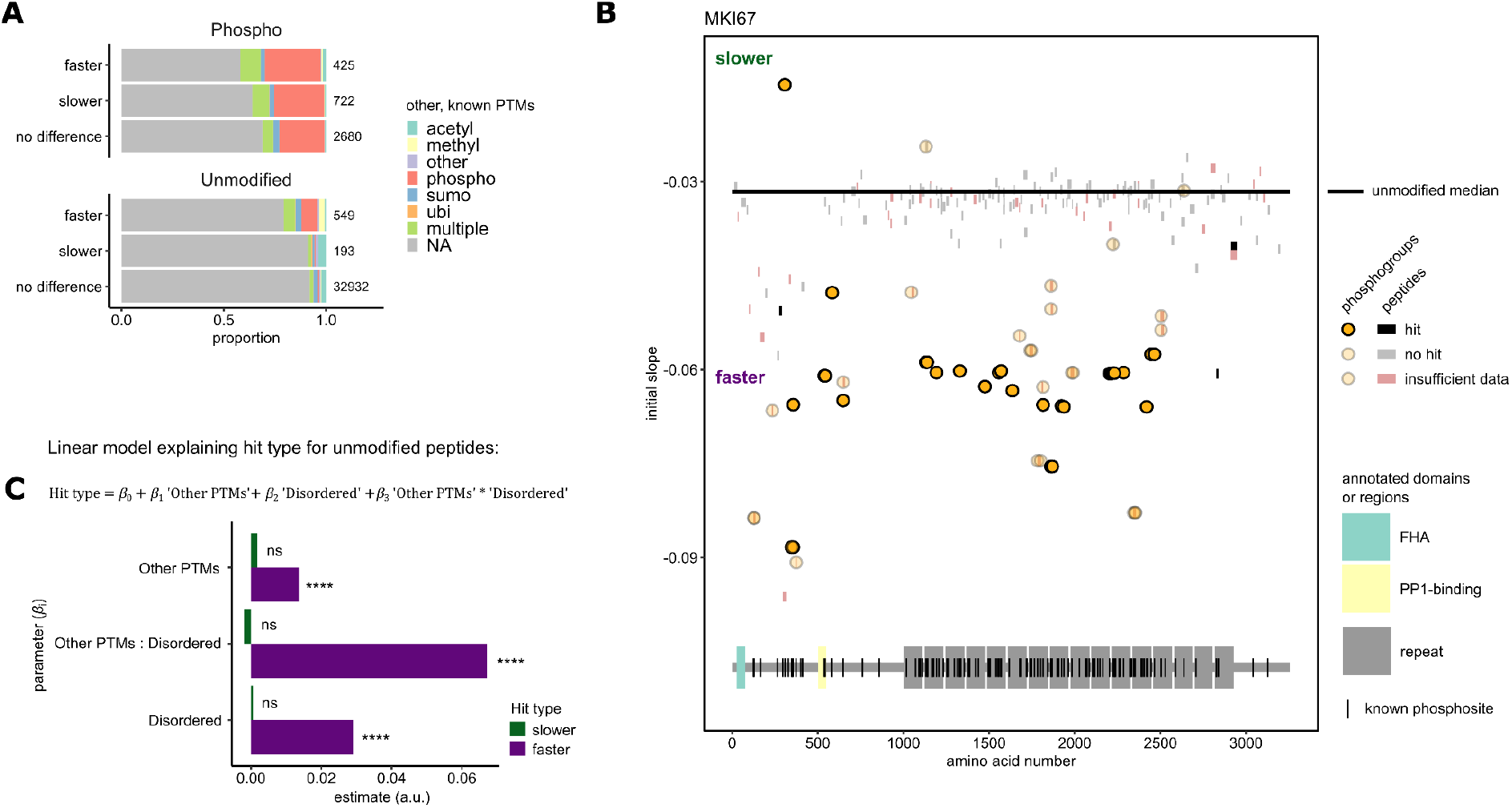
Peptides with faster clearance are enriched in other PTMs and disordered regions. **A**, Distribution of other, known PTMs (from Uniprot) mapping to peptides in our PPToP dataset. **B**, Overview of PPToP data for MKI67 showing substantial number of phosphopeptides with similarly faster clearance. **C**, Peptides with faster clearance are enriched in disordered regions even after correcting for the enrichment of other, known PTMs. Linear regression model used to fit to distribution of slower/faster unmodified peptides. Significance estimated for parameter being non-zero. ****: *p* < 0.0001.

**Figure S5:**
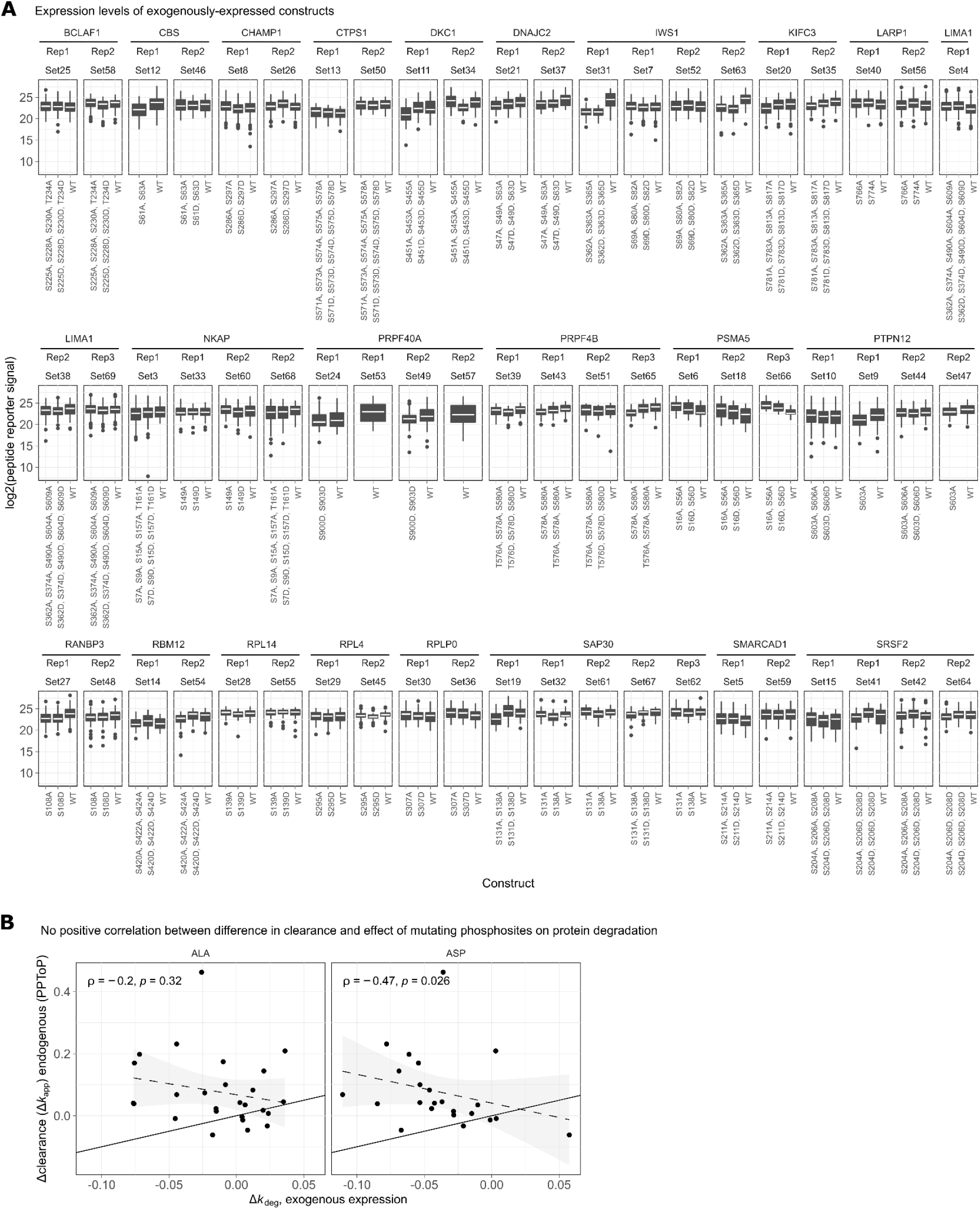
Exogenous expression of GFP-fusion constructs. **A**, TMT reporter signals for LIGHT-labeled peptides of each construct at time-point 0 h of the SILAC time course (see Figure 4 A). Constructs with median expression >5-fold lower than the median of all constructs in that set were excluded from analysis (omitted). Each facet represents data from a single TMT experiment. **B**, No positive correlation between the difference in clearance rate of phosphorylated peptides compared to the protein median (Δ*k*_app_) and the difference in actual degradation constants between mutants of the same phosphorylation sites and the wild-type protein (Δ*k*_deg_).

**Figure S6:**
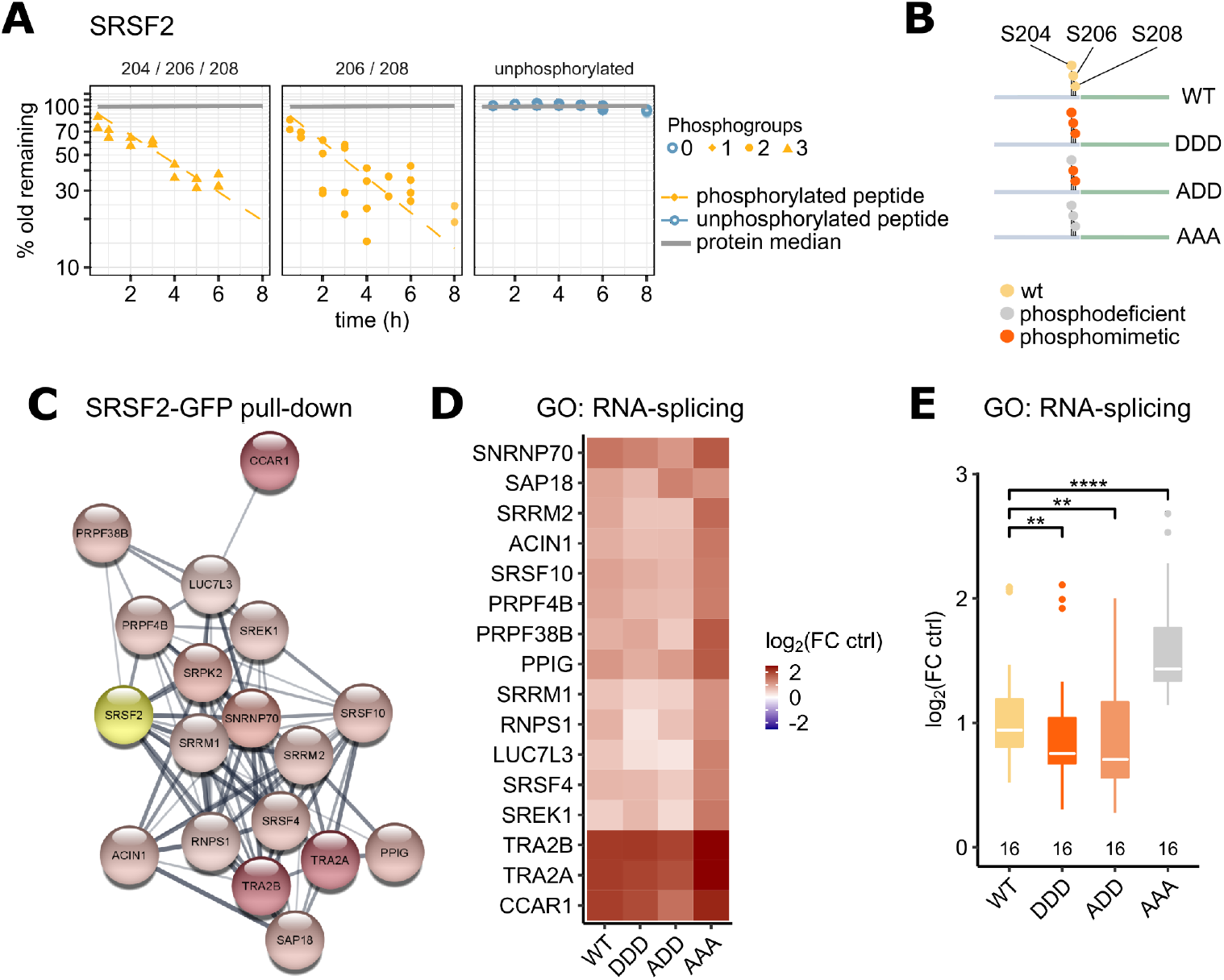
Phosphosites identified by PPToP can define a protein’s protein-protein interaction network. **A**, Faster clearance of phosphopeptides phosphorylated at residues S204, S206, S208. **B**, SRSF2-GFP fusion constructs generated. **C**, STRING interaction network of RNA-splicing proteins pulled-down by SRSF2. Colors represent log2 fold change over control (GFP alone). **D**, Heatmap of same proteins as in (C) for all SRSF2 constructs. **E**, SRSF2 phosphomimicking (ASP, D) mutations show lessened interactions with RNA-splicing factors compared to wild-type, while phosphodeficient (ALA, A) mutations strengthen interaction.

**Figure S7:**
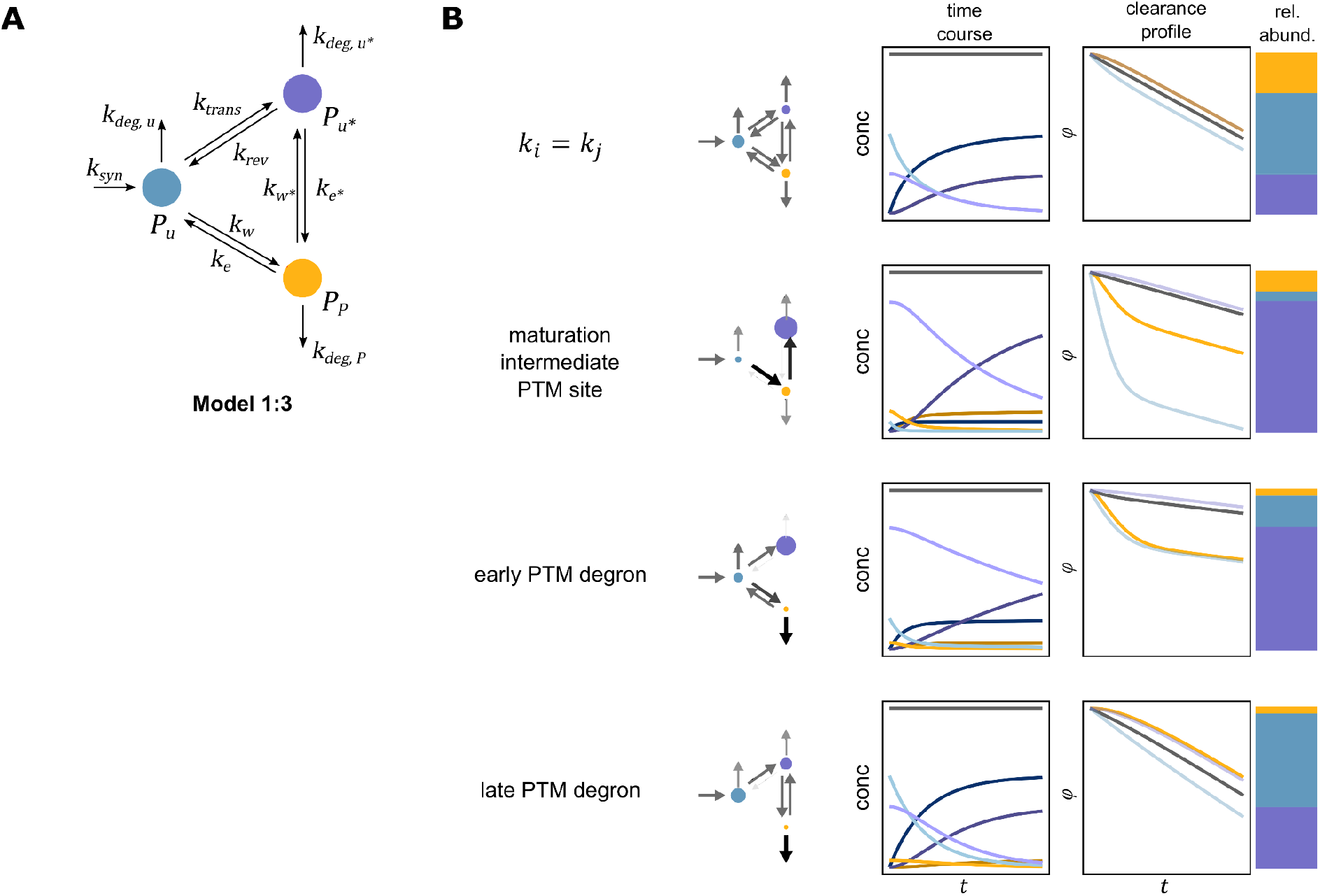
A three-species protein turnover and modification model enables modeling phosphodegrons and intermediate species as species with faster clearance. **A**, Model structure of Model 1:3 with two distinct unmodified species (*P*_u_ and *P*_u*_), which could represent, for instance, young and old protein, or distinct protein pools defined by some other means, such as other PTMs, distinct subcellular localization, protein-protein interaction assemblies, etc. See Theory Supplement for full description of the model. **B**, Example profiles allowed by Model 1:3. First row: all parameters are of the same magnitude. The “early degron” (third row) could, for instance, represent a quality control step early in a protein’s lifetime, where the protein either is modified (and subsequently rapidly degraded), or matures into a more stable form (represented by *P*_u*_). Note that both a maturation intermediate modified pool (second row) as well as an early degron will lead to with faster clearance of the modified species *P*_P_. A “late degron” (bottom row) represents a protein, which only is modified at a later stage during its lifetime. Despite the modified species *P*_P_ having low proteolytic stability (*k*_deg,P_ is high), the clearance profile for *P*_P_ is slower than for the total protein.

**Figure S8:**
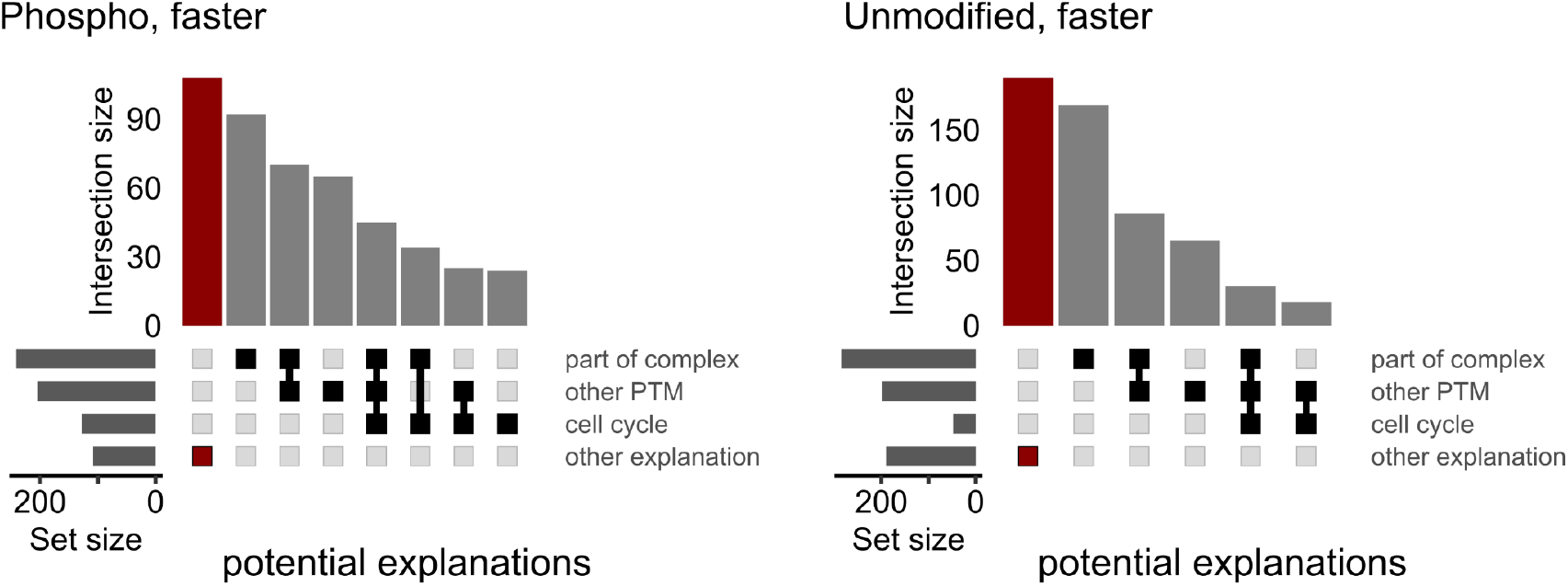
Summary of potential explanations of peptides with faster clearance identified by PPToP. A significant proportion of hits lack a straightforward explanation based on previous data, thus representing an interesting starting point for further discovery.

## Theory Supplement

### 1 Supplement overview

This Supplementary Information contains the theoretical considerations regarding models encoding different wiring schemes of post-translational modifications (PTMs). We use these models to analyse how the clearance profile patterns of Protein-Peptide Turnover Profiling (PPToP) experiments depend on the assumed underlying wiring schemes. Throughout the supplement, we describe and analyse the individual models one by one. The simplest case, where a given protein is modelled as a single, homogeneous pool, has been previously described [Schwanhäusser et al., 2011, Welle et al., 2016]. However, we will use it here to introduce the terminology and illustrate correction of the PPToP data for expansion of the protein pool by cell growth in Section 3.

All symbolic and numeric computations where performed in MATLAB [MATLAB, 2021]. The parameter sample for the plots in Section 4.6 was generated with latin hypercube sampling algorithm from [Khaled, MATLAB Central File Exchange. Retrieved October 05, 2021.] and the plots were generated with the hexscatter function from [Gordon, MATLAB Central File Exchange. Retrieved October 06, 2021.].

### 2 Introduction to the models

The following analysis is based on ordinary differential equations (ODEs) describing the dynamics of a metabolic labelling experiment (Figure 1 A). The ODEs are derived by modelling the wiring scheme based on biochemical reactions obeying mass action kinetics. While we focus mainly on proteoforms defined by the addition and removal of PTMs, the same considerations and conclusions apply to any analogous metabolic labeling experiment of biomolecules in which a species can exist in multiple measurable states that can interconvert, such as nucleic acids and their modifications [Gameiro et al., 2021].

#### 2.1 Modelling assumptions

We built our ODE models on the following assumptions

1. Synthesis is zeroth order.
2. Degradation, as well as addition and removal of a modification is first order and random (i.e. independent of the age of the species in each pool, meaning that the pools are internally homogeneous).
3. The initial conditions correspond to the pre-label-switch steady state.
4. Label switch is near-instantaneous, complete, and label-equilibration times are negligible, as is usually the case in cell culture experiments, where medium exchange is near-total and efficient. Discussion on experimental setups involving slow label equilibration and resulting partial labelling has been provided elsewhere (see, e.g., [Guan et al., 2012, Fornasiero et al., 2018, Hammond et al., 2021]).
5. The label does not affect protein stability or phosphorylation/dephosphorylation kinetics.
6. The data come either from non-dividing cells (constant steady state protein abundance) or have already been corrected for cell cycle-related increase of the protein abundance during data collection. (See Section 3.4 on how to correct for cell cycle effects.)

#### 2.2 Terminology

In the models we refer to protein and peptide species by *P*. No subscript indicates the entire protein, and individual peptide species are indicated by the subscript, e.g. as P*_u_* and P*_P_* for the unmodified and post-translationally modified species, respectively. To distinguish between entities that where synthesized before and after the label switch we append ‘old’ and ‘new’ to the subscript, respectively. The fractional rate constants, that is which fraction of a peptide species is converted per unit time by a reaction, are noted by the letter *k* with a subscript indicating the respective conversion reaction. These rate constants the (unknown) parameters of a model and assumed to be positive. We use 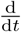 to indicate the derivative of a quantity with respect to time. The definition of a model comprises three components: the ODEs describing the species dynamics, the initial conditions defining the amount of each species at the time of label switch, and the observables encoding the experimental setup by expressing the available data in terms of the model species and parameters. The model name ‘Model x:y’ encodes the number of individual PTM species ‘x’ and the total number of model species ‘y’.

We distinguish between clearance profile and clearance rate to quantitatively describe the readout of PPToP experiments. With *clearance profile* we refer to the time course of the remaining fraction of protein that was synthesized pre-lable switch (old protein). With *clearance rate* we refer to the slope of the time course which describes the speed at which the old protein disappears. Our *clearance rate* therefore corresponds to the *turnover rate* in traditional protein turnover experiments. However, we term it clearance rate to distinguish it from turnover, which is commonly associated with degradation, while we study combined effects of degradation and modification.

### 3 Model 0:1 - a single homogeneous protein pool

In the simplest case a protein is modeled as one homogeneous pool (Figure 1 C). We term this system Model 0:1 (short M0:1, for 0 individual PTM species and 1 protein species in total). In the following subsections we define Model 0:1 by means of its dynamics, initial conditions and observables, derive what we term the clearance profile and clearance rate and recapitulate how to correct for a cell cycle caused increase in protein after the label switch.

#### 3.1 M01: Model definition

Model 0:1 describes the dynamics after the label switch (*t* = 0) as:

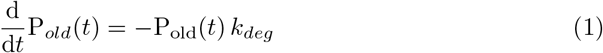

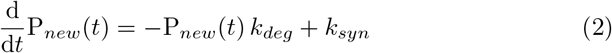

with initial conditions

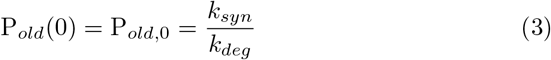

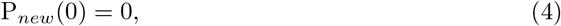

where P_*old*_(*t*) is the number of molecules of protein P that where synthesized prior to the label switch which are present at time *t* after the label switch, P*_new_*(*t*) is the number of molecules of protein P that have been newly synthesized until time *t* after the label switch, *k_syn_* is the synthesis rate, and *k_deg_* the degradation constant of the protein.

The initial condition of the old species, P*_old_*, is determined by the steady state of the model that describes the pre-label-switch system:

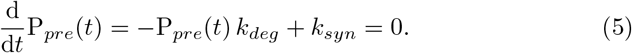

The initial condition of species P*_new_* is 0 because at label switch all protein is old.

The link between a model and the available data is obtained by the definition of model *observables*, which represent the data as a function of the model species. For metabolic labeling experiments, the new protein synthesized after label switch is distinguished from old protein by incorporation of isotopically labelled amino acids. The isotopologue ratio of new and old protein is measured by mass spectrometry, e.g., on the MS1-level Mathieson et al. [2018]. This ratio constitutes the raw data. Accordingly, we can define the observable of Model 0:1 as the fraction of old protein remaining at time *t*, which is derived as follows:

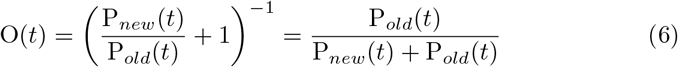

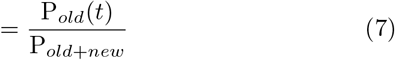

Here the model species ratio 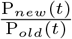 is the model representation of the raw data and we add 1 and take the reciprocal in order to derive the intuitive interpretation as the fraction of old protein remaining. It can be shown that 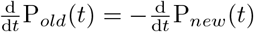. Therefore, the sum of old and new protein is a constant given by the initial abundance of the old protein:

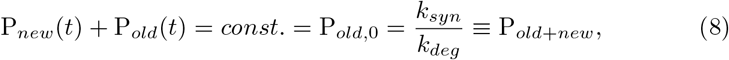

which leads to equation (7).

#### 3.2 M01: Clearance profile and clearance rate

We define the *clearance profile φ*(*t*) as the natural logarithm of the observable (the fraction of old protein remaining at time *t*). For Model M0:1 the clearance profile is given by

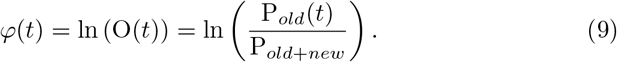

*P*_old_(*t*) can be derived from (1) and (3) to be

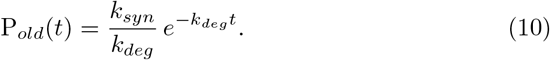

Substituting (10) and (8) in (9), results in the well-known linear relationship

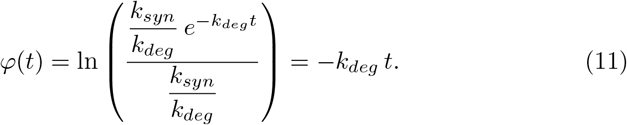

For Model M0:1, the *clearance rate*, which we define as the negative derivative with respect to time (slope) of the clearance profile *φ* (a straight line in case of M01), then recovers the degradation constant from the clearance profile:

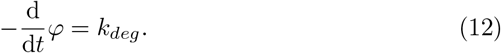

This recovers the well-known fact that the degradation constant of a protein pool can be quantified from the slope of the line through the data on a logarithmic y-axis scale.

#### 3.3 M01: Summary

The clearance profile of Model 0:1 allows to quantify the degradation constant of the protein. Given the profiles of several proteins, it is possible to compare the proteolytic stabilities of those proteins. As shown above, in the case of a protein pool as modelled in Model 0:1, the clearance rate is a constant and identical to the degradation constant *k_deg_*. With this, the degradation constant can quantitatively determined from the slope of the clearance profile. However, the clearance profile as well as the clearance rate are not a function of *k_syn_* and therefore Model 0:1 cannot be used to quantify the rate of protein synthesis.

#### 3.4 A single homogeneous protein pool - Correction for cell cycle related increase in protein abundance

Model M0:1 assumes that the cells are not growing during the data collection. However, in growing cells the total protein amount doubles during the cell cycle, to keep the protein concentration constant. Therefore, in growing cells the sum of old and new protein after the label switch is not constant (as in (8)) but increases as a function of time. For growing cells Model M0:1 is not valid. To describe the dynamics of a homogeneous protein pool in growing cells and to learn how to correct for this effect of the cell cycle we need a slightly different model, described in the following.

In growing cells the dynamics after label switch are described by the following equations

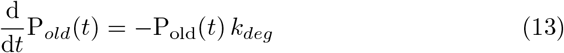

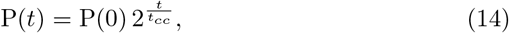

where P*_old_* is equivalent to Model M0:1, P(*t*) is the total protein abundance (old + new) at time time *t* after label switch and *t_cc_* is the cell cycle duration, which is readily measurable in most cell culture experiments, or can be estimated from the data.

Upon label switch, all protein is old, therefore the initial conditions are equal for both model species and given by

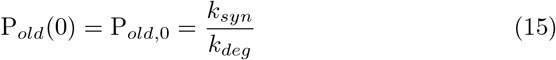

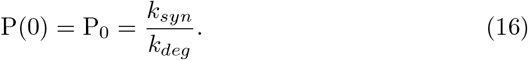

The same reasoning as in Model M0:1 leads to the fraction of old protein remaining as the observable

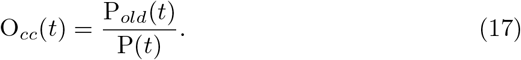

With this the degradation profile including cell cycle effects becomes

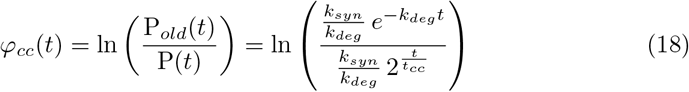

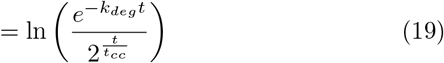

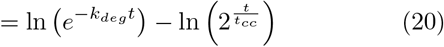

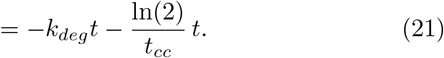

Comparison of (21) and (11), leads to the relation

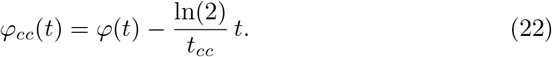

With this, the corrected clearance profile (11) can be derived from an uncorrected clearance profile (21) via the relationship:

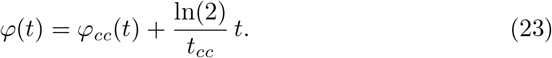

In the same way, 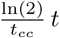 can be used to correct data collected in growing cells at time *t* after the label switch.

### 4 Model 1:2 - synthesis followed by modification

The simplest model for modification of a protein by a PTM consists of two protein species: the unmodified protein P*_u_* and a protein carrying the PTM P*_p_*. Here we assume that a protein is synthesized in an unmodified form and is then reversibly modified in a first order reaction (Figure 1 D). In the following subsections we provide the definition of Model 1:2 (M 12) by means of its dynamics, initial conditions and observables, derive its clearance profile and its clearance rate and provide further analysis of these quantities.

#### 4.1 M12: Model definition

For Model 1:2 the dynamics starting at the time of the label switch (*t* = 0) are given by the ODEs of 4 model species corresponding to two protein species before (old) and after the label switch (new), respectively:

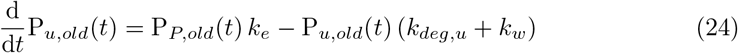

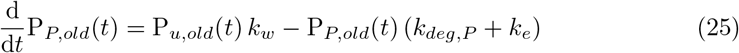

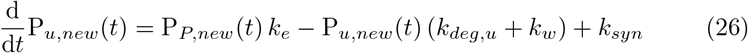

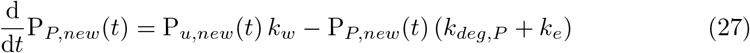

with *k_syn_* being the synthesis rate; *k_deg,u_* the degradation constant of the unmodified species; *k_deg,P_* the degradation constant of the modified species; *k_e_* the rate constant of erasing the PTM; and *k_w_* the rate constant for adding (i.e. writing) the PTM to the protein.

The initial conditions are given as

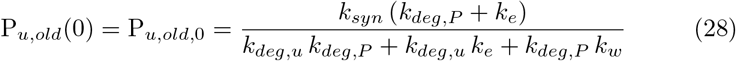

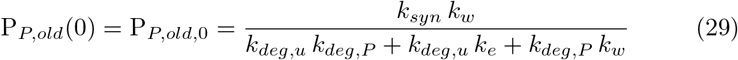

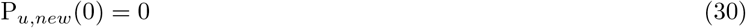

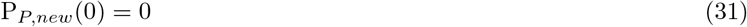

The initial conditions for the old species, P*_u,old_* and P*_P,old_*, are defined by the steady state of the model that describes the pre-switching system:

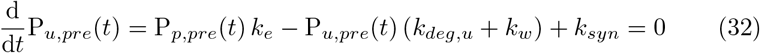

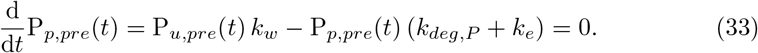

The initial conditions for the new species are 0 because synthesis of the new species only starts at label switch.

Before we go on to define the observables we use some properties of Model 2:1 to reduce the number of equations. This makes numerical as well as symbolical calculations more efficient.

#### 4.2 M12: Model reduction

Although not directly obvious from the model equations (24) to (27), it can be shown that the sum of old and new of each protein species *x* (*x* ≔ {*u, P*}), P*_x,old+new_*, is constant and given by the total abundance of each protein species which is determined by the initial conditions.

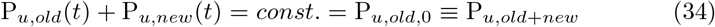

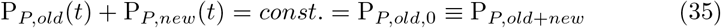

Therefore, the system can be reduced to the following two ODEs:

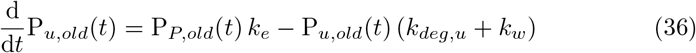

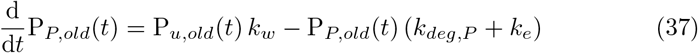

with initial conditions

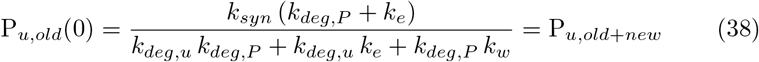

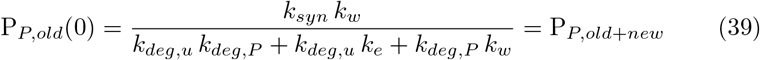

and the two algebraic equations

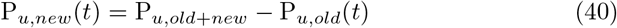

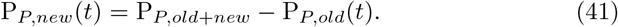

The choice of describing the system using only the subsystem of old species is arbitrary. An equivalent representation could be obtained from the ODEs for the new species (then with initial conditions of zero), and consequently expressing the old species as the difference between the constant total and new. We choose here to focus on the old species since it results in a more intuitive interpretation of the observables, as defined next.

The model observables connect the model species to the available data. For most proteins, the measured data consists of the isotopologue ratios of the modified species, P_*P*_, and an estimate of the total protein, P*_Total_*. P*_P_* is typically defined by a single proteoform-specific peptide, while an estimate of the total protein is given, for instance, by the median of all measured (unmodified) peptides of the protein. As most of the peptides in each protein can be expected to be shared between a modified proteoform (e.g. defined by a single measurable PTM-containing peptide) and the unmodified proteoform, the median will essentially give an abundance-weighted mean estimate over the two proteoforms. It should be noted that in reality many more proteoforms might (and indeed are likely to) exist simultaneously for a given protein, but due to the inherent limitation of bottom-up proteomics, we are in most cases limited to comparing single proteoform-specific peptides to the rest of the peptides.

The measured isotopologue ratios correspond to 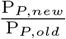 and 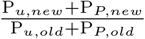 in the model. Equivalently to Model 0:1 we choose to use transformed versions of the data to derive the observables as the fraction of old protein species remaining at each time *t* after label switch. Thus, for Model 1:2, the observables are defined as

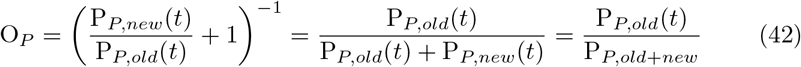

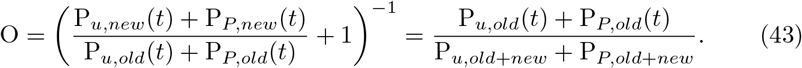

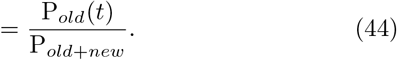

for the entire protein, where P*_old_* refers to total abundance (sum of PTM and unmodified) of old protein left and the constant overall protein abundance is given as a function of the model parameters as

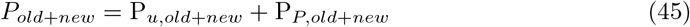

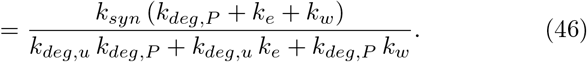

#### 4.3 M12:Analytical solutions of model species time course

The differential equations (36) and (37) can be analytically solved with their initial conditions to yield the time course of the indiviual species

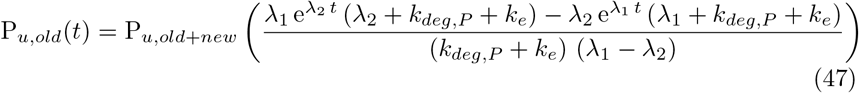

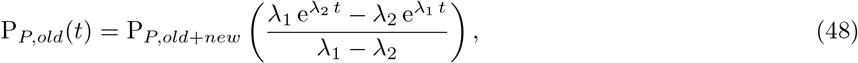

with P*_u,old+new_* and P_*P,old+new*_ as defined in (38) and (39) and λ_1_ and λ_2_ given by

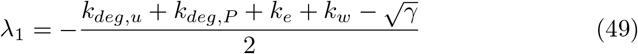

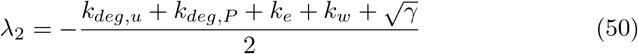

with

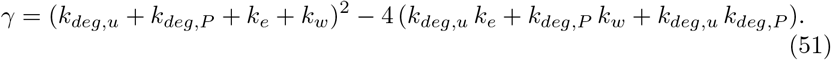

Where λ_1_ and λ_2_ are the eigenvalues of the system matrix

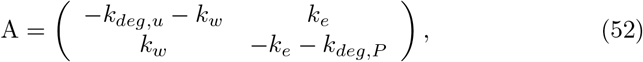

which is derived from the model equations (36) and (37). For Model 1:2 λ_1_ and λ_2_ can only take negative values and λ_1_ will always have a smaller absolute value than λ_2_.

The sum of the two individual species solutions then gives the time course of the remaining old protein

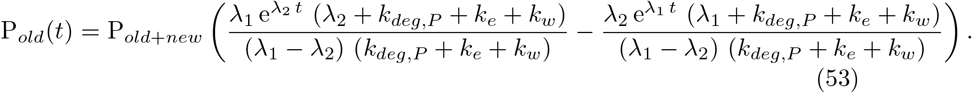

#### 4.4 M12: Clearance profile and clearance rate

With the clearance profiles *φ*(*t*) defined as the natural logarithm of the observables (the fraction of old proteoform remaining at time *t*), the clearance profiles of Model 2:1 become

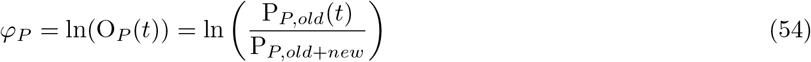

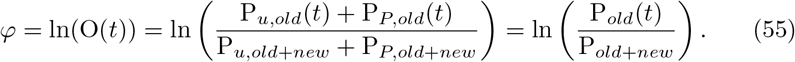

for the PTM species and the entire protein. Note that by dividing the time course function of the PTM species by the total abundance of that species, *k_syn_* cancels out and the profile equation becomes independent of *k_syn_*. The same is true for the profile of the entire protein. Both profiles therefore do not contain any information on *k_syn_*, and *k_syn_* can never be quantified from data.

The clearance rates are then given by the negative of the derivative of the profile with respect to time:

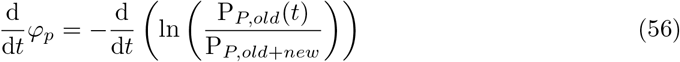

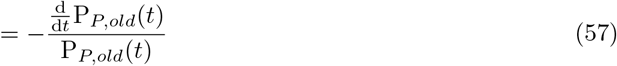

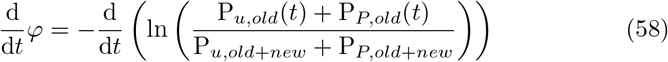

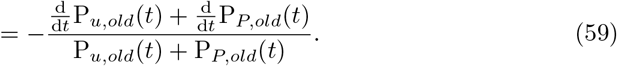

#### 4.5 M12: the PTM clearance rate is always less or equal to the total clearance rate

To show that the clearance rate of the PTM species 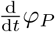 is always less than or equal to the clearance rate of the total protein 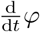, irrespective of the parameter values, we need to show that

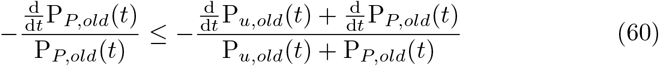

Via rearrangements this leads to

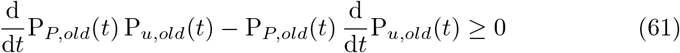

Replacing the derivatives with the model equations and inserting the analytical solutions of the model species results in

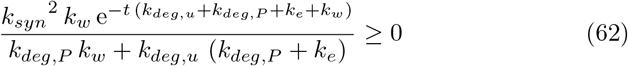

which must be true since we assume all parameters to be positive.

This proofs that the clearance rate of the PTM species is always less or equal to the clearance rate of the entire protein, irrespective of particular parameter values and therefore irrespective of whether *k_deg,P_* is smaller or greater than *k_deg,u_*. Therefore, a comparison between the clearance rates of the PTM species and the entire protein does not contain the information which effect the PTM has on the the protein’s stability. This in in contrast to Model 0:1 where the comparison of the clearance rates of two distinct proteins indicates which of the two proteins is more stable.

#### 4.6 M12: clearance profile curvature as measured by the y-axis intercept of the linear section of the profile

The clearance profiles of the PTM species and the entire protein become linear and parallel at large times after label switch. This indicates that the clearance rates have become equal to the negative of the eigenvalue with the smallest absolute value (see Section 6 for explanation). To quantify the curvature of a clearance profile before it becomes linear, the intercept of the extension of its linear part with the y-axis can be used (see Figure S1 for illustration). With curvature we mean the extend of deviation from of a profile from linearity, e.g. whether a profile is concave up (clearance rate increases over time) or concave down (clearance rate decreases over time) and to what extent.

The y-axis intercept of the linear part of each clearance profile can be derived from the clearance profiles (54) and (55) which leads to

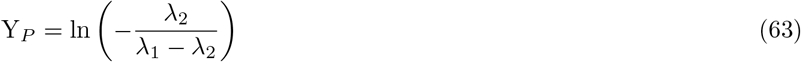

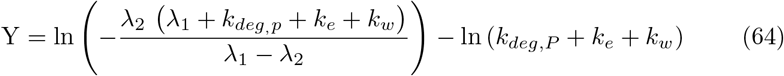

for the intercepts of the linear parts of the PTM and total protein clearance profiles, respectively. Again, λ_1_ and λ_2_ are as given in (49) and (50). Given that λ_1_ and λ_2_ are negative with λ_1_ having the smaller absolute value, Y_*P*_ must always be positive. This is also illustrated in Figure S9 for a set of randomly sampled parameter values. In contrast, the sign of Y depends on which model species has the higher degradation constant. If *k_deg,P_* > *k_deg,u_* (PTM degron) Y can be positive, if *k_deg,P_* < *k_deg,u_* (PTM stabilon) Y can be negative as shown in Figure S10 for the same set of parameters. This corresponds to a concave up shape of *φ* for parameter sets constituting PTM degrons and a concave down shape of *φ* for parameter sets constituting PTM stabilons. Notably, many parameter sets in our random sample result in Y having a value close to 0 (66% for −0.05 ≤ Y*_P_* ≤ 0.05), irrespective whether the parameter set is a PTM degron or a PTM stabilon scenario. This cases correspond to close to linear clearance profiles *φ*.

The difference between the y-axis intercepts of the linear parts of PTM and total protein is given by

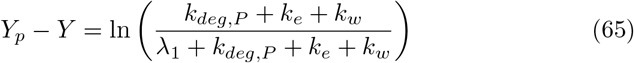

is a quantitative measure of the maximum difference between the clearance rate of PTM and total protein. It must always be positive since λ_1_ is negative and therefore the denominator is smaller than the numerator. Figure S11 illustrates this for a random sample of parameter values. Figure S12 illustrates the relationship between values of the four model parameters and the observed differences between Y*_P_* and Y.

**Figure S9:**
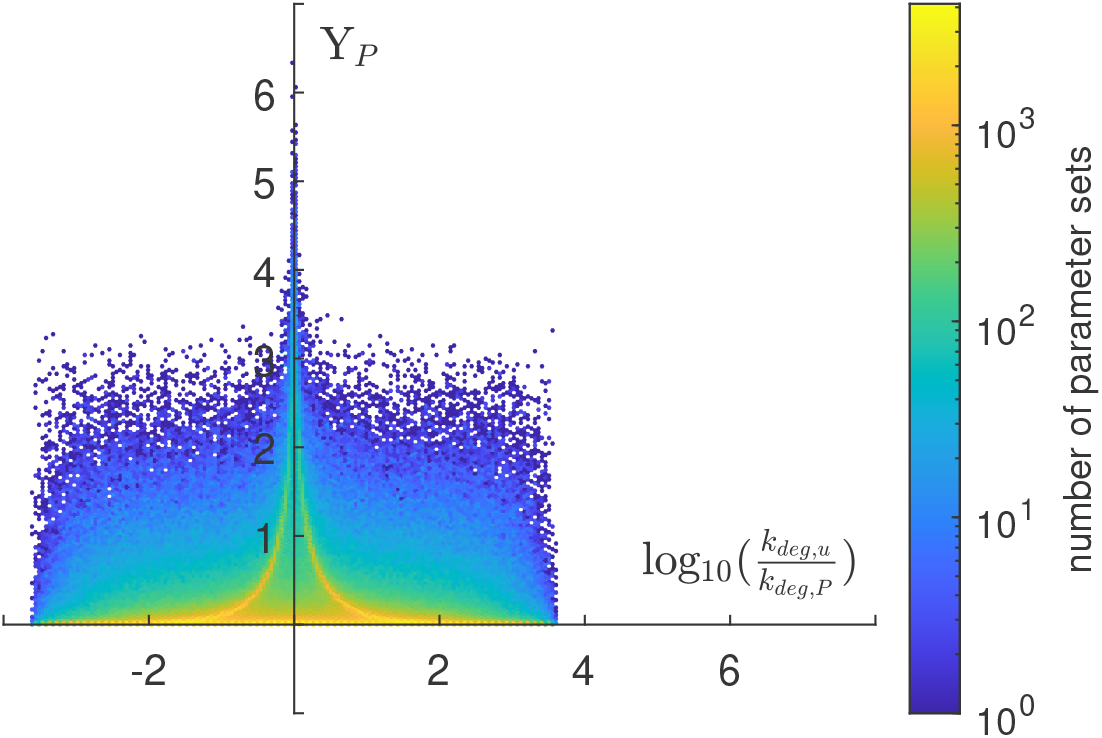
The clearance profile of the PTM species is either linear or concave down. Value of the y-axis intercept of the linear part of the clearance profile of P*_P,old_* (54) versus the logarithm of the ratio between the degradation constants, as calculated for 1 million sets of parameters. Each of the four parameters of Model 1:2 was sampled uniformly on a log2 scale between −12 and 0.

**Figure S10:**
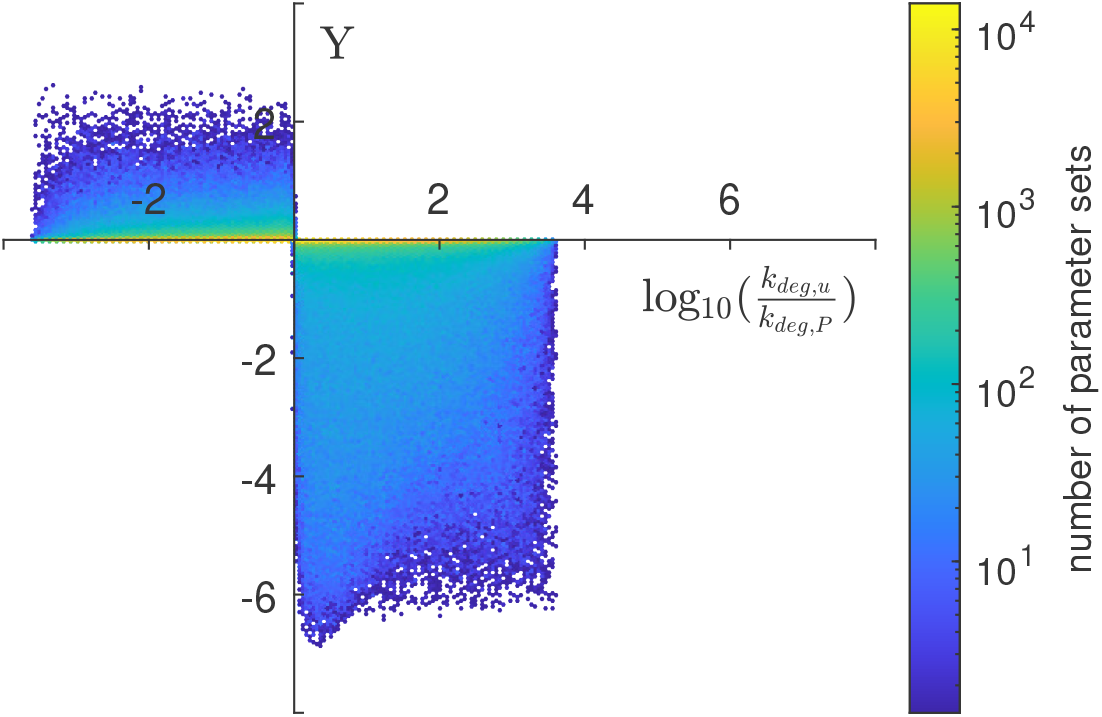
The clearance profile of the entire protein is either linear or concave up (PTM degron) or concave down (PTM stabilon). Value of the y-axis intercept of the linear part of the clearance profile of P*_old_* (55) versus the logarithm of the ratio between the degradation constants, as calculated for 1 million sets of parameters. Each of the four parameters of Model 1:2 was sampled uniformly on a log2 scale between −12 and 0.

**Figure S11:**
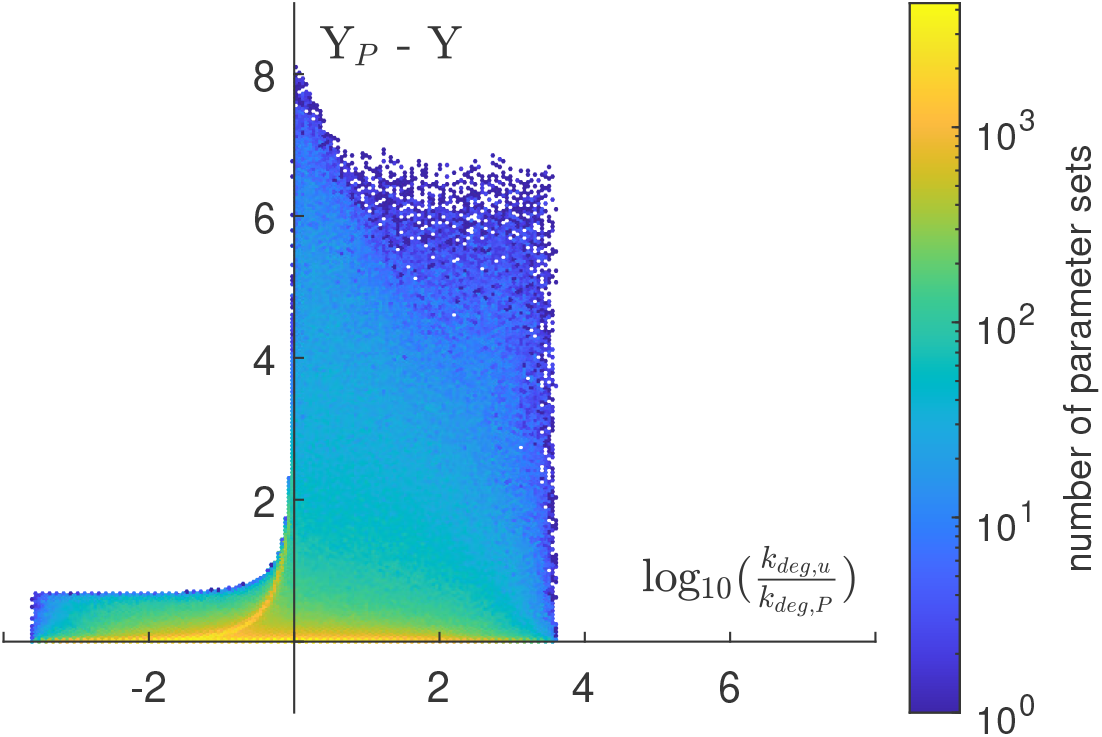
The distance between the profiles of PTM species and entire protein is always positive. Difference between the y-axis intercepts of the linear parts of the clearance profiles versus the logarithm of the ratio between the degradation constants, as calculated for 1 million random sets of parameters. Each of the four parameters of Model 1:2 was sampled uniformly on a log2 scale between −12 and 0.

#### 4.7 M12: Structural identifiability analysis

In the same way that model M01 is frequently used to quantify the degradation constant (or half life) of protein pools, models could be used to estimate the model parameters from experimental clearance profile data. Quantitative values for the parameters could then be used to determine the effect of a PTM on protein stability, i.e. to call PTM degrons and PTM stabilons. The first step in the process of parameter estimation should always be a structural identifiability analysis of the model [Villaverde et al., 2022].

Structural identifiability analysis answers the question whether it is possible to quantify the unknown parameters of a model by means of an analysis solely considering the model structure, i.e. the model equations together with the initial conditions and the observables. Since the analysis does not consider any data it is also called *a priori* identifiability analysis. Structural identifiability is a binary model property - a model is either identifiable or not. A model is structurally identifiable if all of the unknown parameters are structurally identifiable. If one or more parameters are not structurally identifiable (structurally unidentifiable) the model is not structurally identifiable. Structural identifiability is a prerequisite for the quantification of a parameter and its uncertainty from real data. However, how well a parameter can be quantified, ultimately depends on the quality and quantity of the data.

There are multiple methods to perform a structural identifiability analysis, some of which have a readily available implementation [Chis et al., 2011]. The user provides the model structure (model equations including inputs + initial conditions+ observables) and the analysis tool returns whether the provided model structure is structurally identifiable, usually together with information on the identifiability of the individual parameters. We used DAISY [Bellu et al., 2007] and STRIKE-GOLDD [Villaverde et al., 2016] to analyse Model 1:2. The analysis revealed that out of the five model parameters only *k_deg,u_* is structurally identifiable.

**Figure S12:**
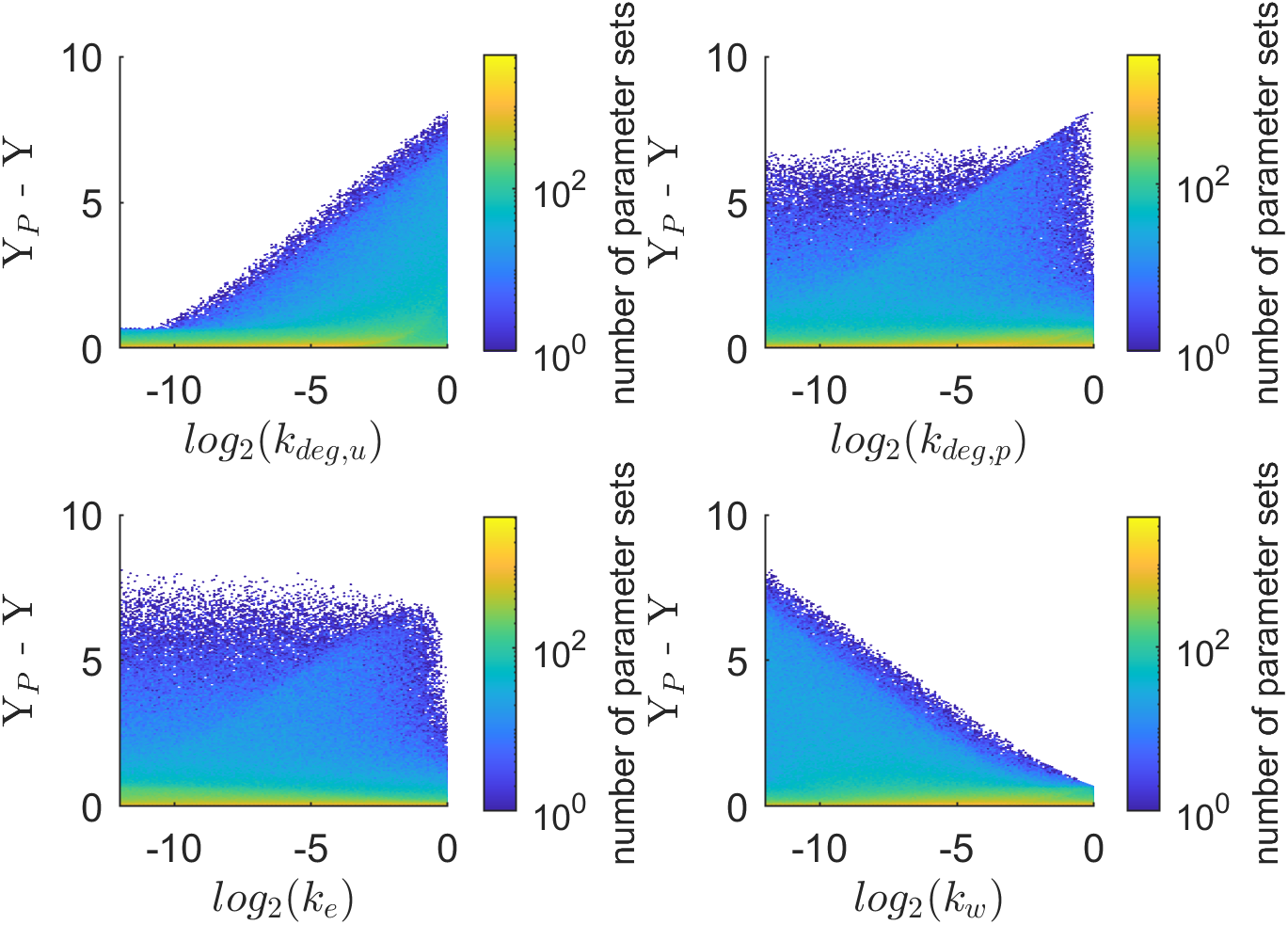
A PTM species profile that is much slower than the profile of the entire protein is caused by a big *k_deg,u_* in combination with a small *k_w_*. Difference between the y-axis intercepts of the linear parts of the clearance profiles versus log2 of the value of each of the four parameters in each of the 1 million random sets of parameters.

We analysed several variants of Model 1:2 that differ in the model observables. In addition to the observable defined in (82) and (85) we consider the fraction of old unmodified protein remaining as a possible observable:

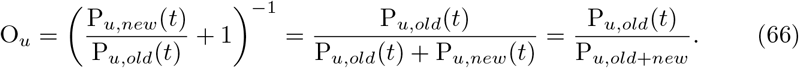

Table 1 summarises the the identifiability properties of the model variants defined by the set of observables. In summary, *k_syn_* is unidentifiable for all three theoretically possible observable combinations. This is expected since the observable are not a function of *k_syn_*. To make all other parameters identifiable one would need to measure the remaining fraction of the old unmodified species in addition to the remaining fraction of the total old protein. Notably, obtaining the clearance profiles of the two most readily-measurable species (the modified species P*_P_*, which can be enriched with specific PTM-enrichment, and the total P, which can be estimated from all unmodified peptides of the protein) only allows for structural identifiability of *k_deg,u_*. Furthermore, as the unmodified species P*_u_* is only biologically meaningful in conjunction with its designated modified counterpart, measuring it alone (while being the most information rich for the wiring scheme encoded in Model 1:2 from the standpoint of identifiability) wouldn’t readily yield biological insight.

**Table 1:** Observables in different variants of Model 1:2 and structurally identifiable parameters are indicated by “x” in the respective column. The first row corresponds to Model 1:2.

| O | $O_P$ | $O_u$ | $k_{syn}$ | $k_{deg,u}$ | $k_{deg,P}$ | $k_e$ | $k_w$ |
| --- | --- | --- | --- | --- | --- | --- | --- |
| x | x |  |  | x |  |  |  |
|  | x | x |  |  |  |  |  |
| x |  | x |  | x | x | x | x |

#### 4.8 M12: Summary

For Model 1:2 the order of the clearance profiles (slower/faster) contains no information about the proteolytic stability (*k_deg,x_*) of the different species. The profile of the modified species is always slower then the profile of the entire protein. The clearance rates are functions of time. The clearance rate of the modified species starts at 0 and increases until it becomes constant at λ_1_, The clearance rate of the entire protein starts at a non-zero value and decreases or increases (depending on whether its a PTM stabilon or PTM degron) until it reaches λ_1_.

### 5 Model 1:3 - alternative path to modification via an additional protein species

Adding one more unmodified proteoform to the mechanism, we can derive a general three-species model consisting of two unmodified species and one PTM protein species (Figure S7). We assume that a protein is synthesized in an unmodified form, P_*u*2_, and then this species is either directly modified as in Model 1:2, or can access another unmodified state P_*u*2_. Here, P_*u*2_ corresponds to a distinct protein pool defined by some other means, such as other PTMs, distinct subcellular localization, protein-protein interaction assemblies, or some other feature, which would make it behave in a distinct manner from the other unmodified proteoform P_*u*1_.

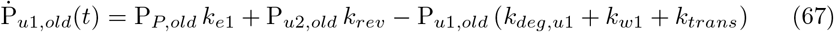

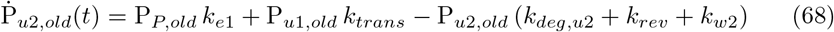

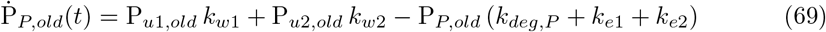

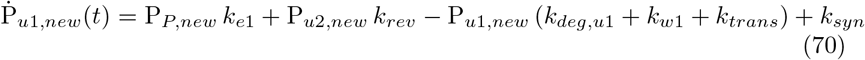

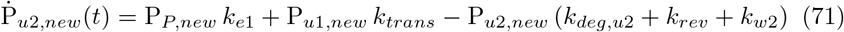

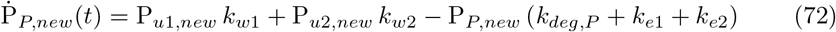

where each of the protein species has its individual degradation constant *k_deg,x_ x* = {*u*1, *u*2, *p*}, and the remaining rate constants describe the kinetics of interchange between the different protein forms.

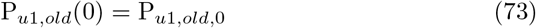

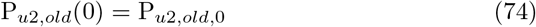

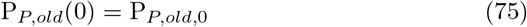

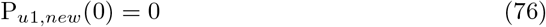

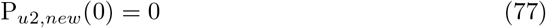

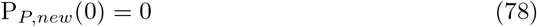

The initial conditions for the old species, P_*u*1,*old*_, P_*u*2,*old*_ and P*_P,old_*, are defined by the steady state of the model that describes the pre-switching system:

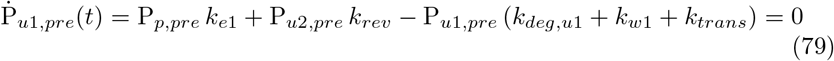

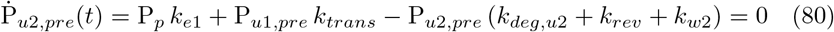

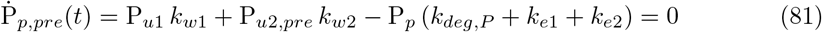

while the ones for the new species are 0 since all protein is old.

The measured isotopologue ratios correspond to 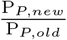 and 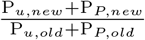 in the model. Equivalently to Model M0:1 we choose to use transformed versions of the data to derive the observables as the fraction of old protein species remaining at each time *t* after label switch. Thus, for Model 1:3, the observables are defined as

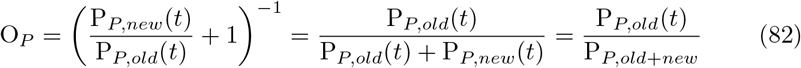

for the PTM proteoform and

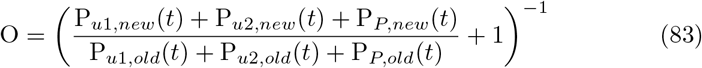

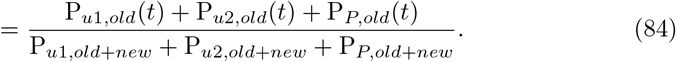

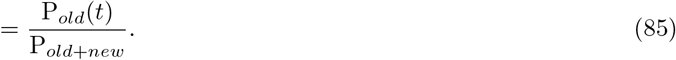

for the total old protein abundance, where P_*old*_ refers to total abundance (sum of PTM and unmodified) of old protein left and the constant overall protein abundance is given as a function of the model parameters as

### 6 General analysis of clearance profiles and clearance rates

We define the *clearance profile* as the natural logarithm of the fraction of old protein of species P*_x_* remaining, *φ_x_*(*t*), at time *t*, for any protein species P*_x_* as:

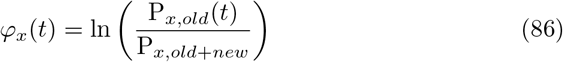

where the *x* in the subscript is used to refer to the different model species in case there is more than one.

Accordingly, we define the *clearance rate* as the negative of the derivative of the clearance profile with respect to time (the slope) as

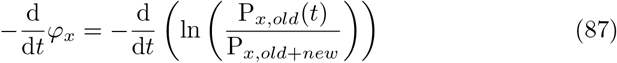

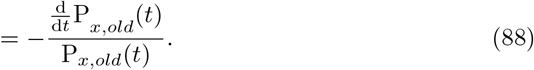

We take the negative for the clearance rate to be a positive number.

To assess the clearance rate we thus we need to look the analytical solution of the protein species P*_x,old_*(*t*) and its derivative only. For linear differential equations it is often possible to derive this solution, which is in its general form given by

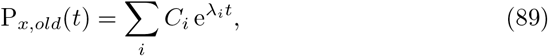

where *i* is the number of protein species involved in the mechanism (i.e. two for Model 1:2), λ_*i*_ is the *ith* eigenvalue of the system matrix *A*, which is derived from the model equations. The eigenvalues are functions of the parameters, including the degradation constants of each species. The coefficients *C_i_* are constants which are functions of the parameters.

The derivative of this solution with respect to time can then be obtained to be

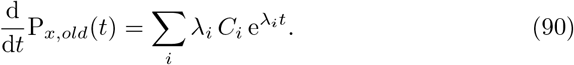

Using this general representation of the solution and its derivative, the clearance profile can be written as

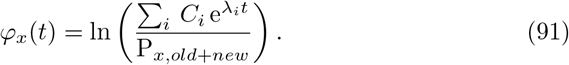

Correspondingly, the general representation of the clearance rate is given by

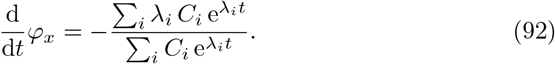

Equation (92) shows that the clearance rate is not just a function of time, but the relation between the clearance rate and the degradation constant becomes more complicated, the more species are involved in a wiring scheme.

If we order the eigenvalues by increasing absolute value, the exponential that has λ_1_ in the exponent will be the exponential that declines to 0 the slowest. Therefore, this exponential function be the sole exponential left for large times *t*:

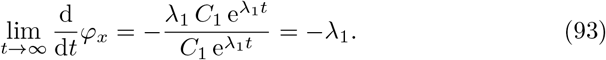

Hence, for large times the clearance profile becomes linear with the clearance rate (negative slope) given by λ_1_.

